# Card9 and MyD88 differentially regulate Th17 immunity to the commensal yeast *Malassezia* in the murine skin

**DOI:** 10.1101/2024.07.12.603211

**Authors:** Meret Tuor, Mark H.T. Stappers, Fiorella Ruchti, Alice Desgardin, Florian Sparber, Selinda J. Orr, Neil A.R. Gow, Salomé LeibundGut-Landmann

## Abstract

The fungal community of the skin microbiome is dominated by a single genus, *Malassezia*. Besides its symbiotic lifestyle at the host interface, this commensal yeast has also been associated with diverse inflammatory skin diseases in humans and pet animals. Stable colonization is maintained by antifungal type 17 immunity. The mechanisms driving Th17 responses to *Malassezia* remain, however, unclear. Here, we show that the C-type lectin receptors Mincle, Dectin-1, and Dectin-2 recognize conserved patterns in the cell wall of *Malassezia* and induce dendritic cell activation *in vitro*, while only Dectin-2 is required for Th17 activation during experimental skin colonization *in vivo.* In contrast, Toll-like receptor recognition was redundant in this context. Instead, inflammatory IL-1 family cytokines signaling via MyD88 were also implicated in Th17 activation in a T cell-intrinsic manner. Taken together, we characterized the pathways contributing to protective immunity against the most abundant member of the skin mycobiome. This knowledge contributes to the understanding of barrier immunity and its regulation by commensals and is relevant considering how aberrant immune responses are associated with severe skin pathologies.

## INTRODUCTION

The skin is the largest organ of the human body providing a complex interface for microbial interactions. It is an important physical and immunological barrier for protection of the host from environmental insults to which it is constantly exposed, such as mechanical damage, toxic chemicals, and pathogenic infectious agents. Like other epithelial barrier tissues, the skin is populated by a wide variety of commensal microbes including bacteria and fungi that contribute to tissue homeostasis and host physiology [1]. While commensals exhibit host-beneficial properties, it is essential to tightly regulate their growth to prevent the development of host-adverse activities, potentially resulting in pathogenicity. The skin mycobiome is dominated by basidiomycetous yeasts of the genus *Malassezia* [2]. Twenty *Malassezia* species have been identified to date [3]; *M. pachydermatis* is commonly found on warm-blooded animals, especially dogs and cats. *M. restricta* and *M. globosa* are most abundant on human skin, followed by *M. sympodialis* and *M. furfur* [3]. Although found in all areas of the skin, *Malassezia* spp. are enriched in sebaceous sites, which is self-evident given their dependence on exogenous lipid sources for thriving, due to the lack of fatty acid synthase [4]. Hair follicles, for example, represent a preferred niche for the lipophilic yeasts as they provide secreted host lipids as a nutrient source [5]. Given the pathogenic potential of *Malassezia* due to host (genetic) predisposition, dysbiosis or other conditions [6], [7], [8], tight immunological control is needed to maintain homeostasis.

The immune system plays a pivotal role in maintaining stable host-fungus interactions in barrier tissues. Protective immunity against *Malassezia* depends on interleukin-17 (IL-17)- mediated immunity [9] and healthy individuals bear *Malassezia*-responsive memory T helper 17 (Th17) cells that readily produce the cytokines IL-17A and IL-17F, i.e. the main representatives of the IL-17 cytokine family [10], [11]. The prevalence of seborrheic dermatitis, an inflammatory skin disorder associated with fungal overgrowth, is enhanced in HIV^+^ individuals bearing low CD4^+^ T cell counts[12]. Complementary to these findings in humans, it has been shown that experimentally colonized mice also mount a strong type 17 immune response that is mediated by αβ and γδ T cells directed against *Malassezia* [10], [13]. Accordingly, T-cell-deficient mice and mice lacking a functional IL-17 pathway are unable to prevent fungal overgrowth [10]. While the relevance of type 17 immunity for immunosurveillance of *Malassezia* commensalism has been established, it remains less clear, how protective Th17 cells are induced in response to *Malassezia*. T cell priming in response to microbes relies on the capacity of the host to sense *Malassezia* spp. via pattern recognition receptors (PRRs). PRRs excel by discriminating different classes of microbes [14]. C-type lectin receptors (CLRs) are of particular importance for sensing fungi, given the specificity of many CLRs for carbohydrate moieties that are abundant in fungal cell walls [15], [16] and for coupling fungal recognition to adaptive immune activation via a pathway that involves the kinase Syk and the adaptor Card9 [17]. Although little studied, the cell wall of *Malassezia* has been shown to differ markedly from that of other fungal genera by its high chitin and chitosan content and the abundance of 1,6-linked β-glucan [18]. How this cell wall is sensed by the immune system remains incompletely understood. CLRs and TLRs have been reported to be involved [19], [20], [21], but the role of these pathways, the relative contribution of individual receptors and how they link innate and adaptive antifungal immune responses in the *Malassezia*-colonized skin *in vivo* has not been elucidated.

Here we comprehensively and comparatively dissected the role of CLRs, Toll-like receptors (TLRs), and IL-1 family cytokines in *Malassezia*-specific Th17 immunity in the skin - the natural habitat of the fungus. Using an experimental model of epicutaneous fungal colonization in mice, we found that Dectin-2 is required for coupling innate fungal sensing to the induction of cutaneous Th17 immunity against *Malassezia*, while Mincle and Dectin-1 were redundant. We also performed an in-depth characterization of myeloid cell dynamics including dendritic cells (DCs) in the ear skin and skin draining lymph nodes (dLN) upon *Malassezia* colonization, which revealed that the conventional DC2 (cDC2) subset of DCs is specifically activated while the cDC1 subset and Langerhans cells (LC) are redundant for efficient priming of *Malassezia*-specific Th17 cells. Furthermore, this study shows that MyD88-dependency of the antifungal Th17 response is attributed to IL-1 family cytokine signaling.

## RESULTS

### The C-type lectin receptors Mincle, Dectin-1, and Dectin-2 bind to *Malassezia* spp

To understand which PRRs may be involved in innate immune recognition of *Malassezia*, we explored a transcriptomic dataset of *M. pachydermatis* -colonized murine skin [22]. We checked for differential expression of PRR encoding genes in colonized vs. naïve skin using the GO term “pattern recognition receptor activation”, GO:0038187. Among the most strongly upregulated genes were those encoding the CLRs Mincle (*Clec4e*), Dectin-3 (*Clec4d*), Dectin-1 (*Clec7a*) and Dectin-2 (*Clec4n*) (**Fig. 1A**). Considering that Dectin-3 primarily acts in association with other CLRs [23], [24], [25] we focused on Mincle, Dectin-1 and Dectin-2 ^19–21^. To test the binding capacity of these receptors to *Malassezia*, we made use of soluble murine CLR-Fc constructs [26], [27], [28]. Mincle, Dectin-1 and Dectin-2 all specifically bound to live fungal cells of three distinct *Malassezia* species: *M. pachydermatis, M. sympodialis.* and *M. furfur* (**Fig. 1B-D**, **Fig. S1A**). Visualization of the bound CLR-Fcs by microscopy revealed a strong signal at the fungal cell surface (**Fig. 1B**), in line with the CLRs binding to fungal cell wall components. Quantification of the binding by flow cytometry confirmed the specific interaction of the receptors with *Malassezia* spp. when compared to a non-relevant Fc construct (CR-Fc in case of Dectin-1 and Dectin-2) or to a control staining with the secondary detection antibody only (2nd (amIgG) in case of Mincle) The signal for either control was as low as a completely unstained sample (**Fig. 1B**). For each of the receptors, the binding strength was comparable across all three tested fungal species (**Fig. 1C-D**). Together, these data suggest that in line with previous reports [19], [20], [21], Mincle, Dectin-2 and Dectin-1 served as receptors for the skin commensal yeast and that the molecular patterns recognized by these receptors are conserved across *Malassezia* species.

**Figure 1.**
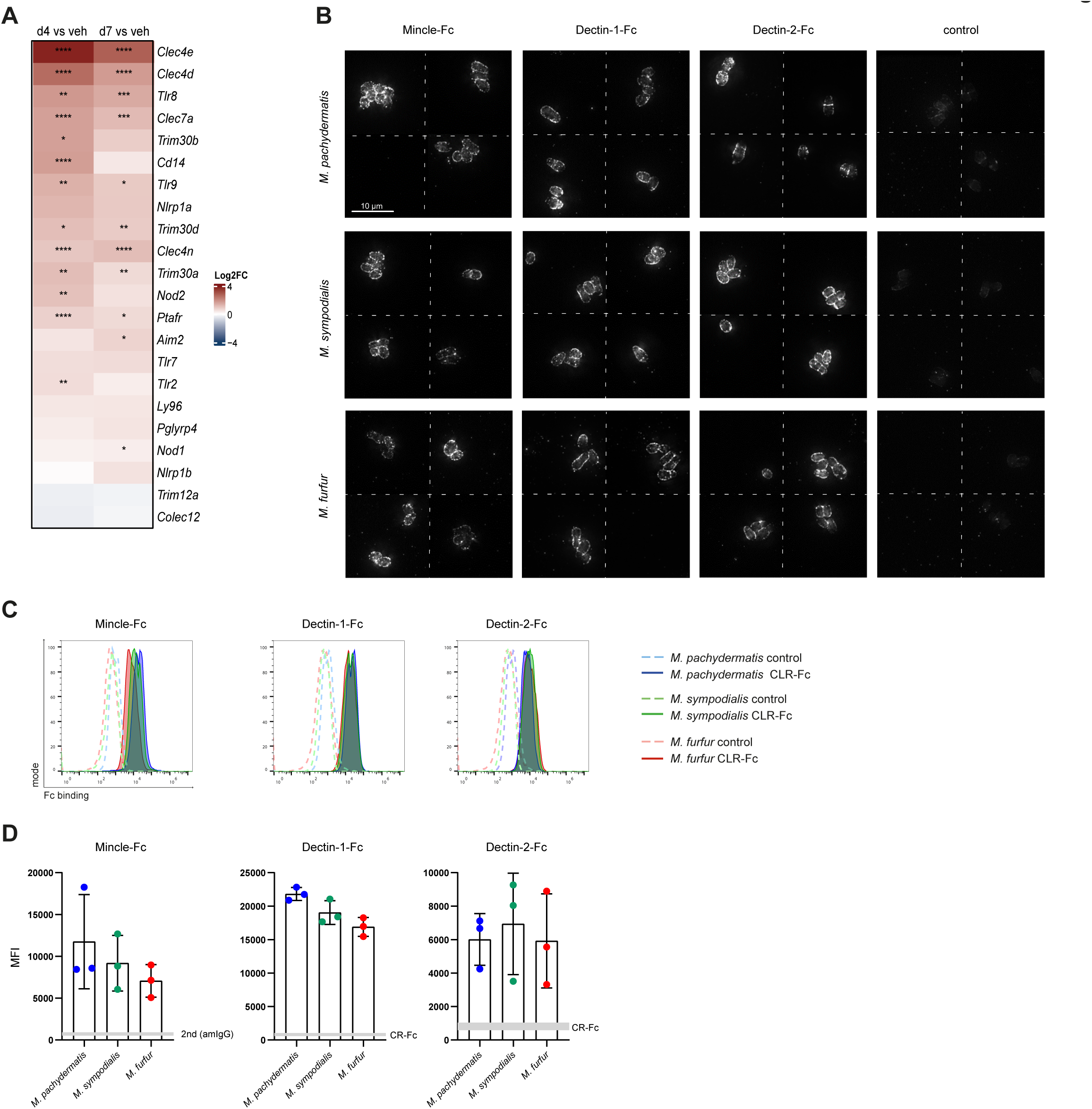
The C-type lectin receptors Mincle, Dectin-1, and Dectin-2 bind to *Malassezia* spp. **A.** Differentially regulated genes (log2 fold change) linked to GO: 0038187 “Pattern Recognition Receptor Activity” in the ear skin of mice colonized for 4 days (d4) or 7 days (d7) with *M. pachydermatis* in comparison to vehicle-treated mice (veh). **B-D.** Live *M. pachydermatis*, *M. sympodialis* and *M. furfur* cells were incubated with Mincle-Fc, Dectin-1- Fc, Dectin-2-Fc or CR-Fc (control) and analyzed by microscopy (B) or flow cytometry (C-D). 4 representative images are shown for each receptor and control staining condition are shown in B. The scale bar represents 10 μm. Representative histograms and the median fluorescence intensity (MFI) of CLR-Fc binding are shown in C and D, respectively. Data are from one experiment (B), from one representative of 3 independent experiments (C), or pooled from three independent experiments with n=1 each (D). **See also Fig. S1**.

### CLR-Card9-dependent signaling in response to *Malassezia* activates dendritic cells

To assess whether the engagement of CLRs by *Malassezia* translates into downstream signaling and cellular activation, we examined cytokine production by DCs upon *Malassezia* exposure. DCs are among the immune cell types that express CLRs most strongly, and they play a central role in coupling microbial recognition to T cell activation [17], [29], [30]. We first broadly assessed the involvement of CLR signaling during *Malassezia*-induced activation of DCs by testing the dependence on Card9, i.e. the common downstream signaling adapter of most CLRs [31]. For this, we stimulated bone marrow-derived dendritic cells (BMDCs) from Card9-sufficient and -deficient mice with *Malassezia* spp. for 24 h. The strong induction of IL-12/23p40 secretion by live *M. pachydermatis*, *M. sympodialis*, and *M. furfur* was strongly Card9-dependent (**Fig. 2A**). Stimulation with CpG confirmed that *Card9*^-/-^ BMDCs were fully competent to produce cytokines via other PRR pathways, while the response to the β-1,3-glucan curdlan, a specific Dectin-1 agonist, was also abolished in absence of Card9 (**Fig. S2A**), as expected [17]. Cytokine levels in uninfected controls were as low as the detection limit. Expression of the IL-23p19 receptor subunit (*Il23a*) and of IL-6 was also strongly induced by *Malassezia* spp. in a Card9-dependent manner (**Fig. 2B, Fig. S2B-C**), both being implicated in Th17 polarization. To determine the impact of individual CLRs on DC activation, we next tested the consequence of Mincle-, Dectin-1- or Dectin-2-deficiency on *Malassezia* recognition and DC activation. Deficiency in either of the receptors led to significant reduction of cytokine secretion in response to *M. pachydermatis*, *M. sympodialis*, and *M. furfur* (**Fig. 2C-E**), while competency of the gene deficient BMDCs to Card9- independent stimulation was confirmed (**Fig. S2D-F**). Because cytokine responses were only partially reduced in Mincle, Dectin-1, and Dectin-2 deficient BMDCs in contrast to the almost complete impairment in Card9 deficient cells, we speculated that individual CLRs may display functional redundancy, at least in part, for *Malassezia*-induced cytokine response by DCs. We therefore assessed the responsiveness of BMDCs lacking both, Dectin-1 and Dectin-2 (Dectin-1-Dectin-2 double knockout, (DKO)) or BMDCs deficient in all three aforementioned CLRs Mincle, Dectin-1 and -2 (Mincle-Dectin-2-Dectin-1 triple knockout, (TKO)) [32]. Cytokine levels of DKO BMDCs were reduced to baseline levels in response to *Malassezia* spp., similarly to what was observed for Card9-deficient BMDCs (**Fig. 2F, Fig. S2G**). Additional deletion of Mincle did not further reduce the response (**Fig. 2F, Fig. S2G**). This highlights that Dectin-1 and Dectin-2 collectively mediate recognition of *Malassezia* by DCs, suggesting a pivotal role of these two CLRs for mounting protective antifungal responses in the colonized skin.

**Figure 2.**
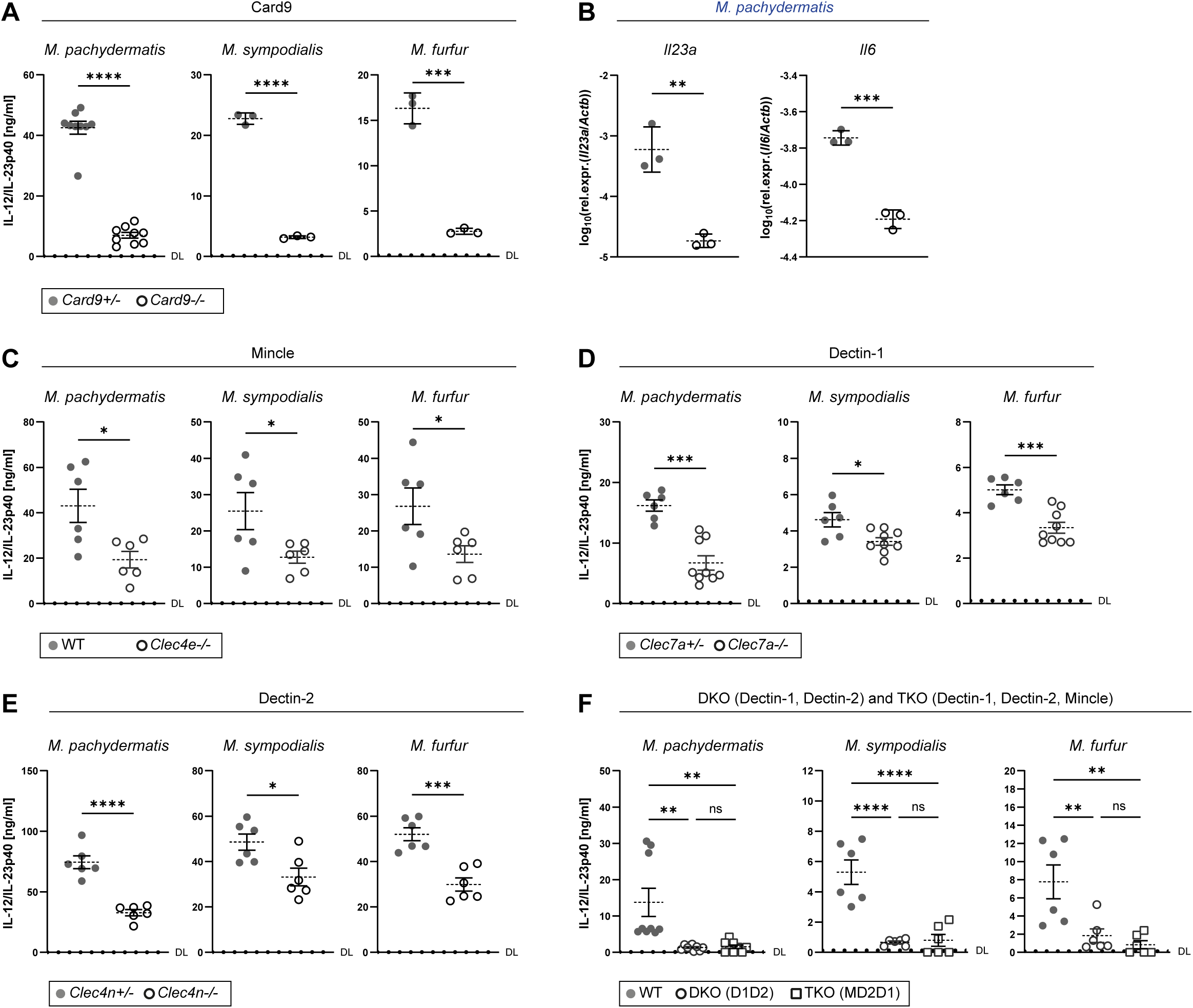
CLR-Card9-dependent signaling in response to *Malassezia* activates dendritic cells. **A-F.** BMDCs generated from *Card9*^+/-^ and *Card9*^-/-^ mice (A-B), WT control mice and *Clec4e*^-/-^ (C), *Clec7a*^+/-^ and *Clec7a* ^-/-^ mice (D), *Clec4n*^+/-^ and *Clec4n* ^-/-^ mice (E), and WT control mice, Dectin-1-Dectin-2 DKO, and Mincle-Dectin-2-Dectin-1 TKO mice (F) were stimulated with live *M. pachydermatis*, *M. sympodialis* or *M. furfur* cells for 24 h. BMDC activation was assessed by sandwich ELISA for IL-12/IL-23p40 protein (A, C-F) and by RT-qPCR for *Il23a* and *Il6* transcripts (B). Each symbol represents a separately stimulated well. The mean ± SEM is indicated for each group and data are pooled from two to three independent experiments with n = 3 per group each. Data in A (*M. sympodialis* and *M. furfur*) and B are the mean ± SD from one experiment. DL = detection limit. Statistical significance was determined using unpaired t test (A-E) or one-way ANOVA (**F**), * p < 0.05, ** p < 0.01, *** p<0.001, **** <0.0001. **See also Fig. S2**.

### *Malassezia* skin colonization recruits myeloid cells and activates *cDC2s*

To study CLR-mediated immune activation by *Malassezia in vivo*, we used an experimental model of skin colonization using *M. pachydermatis* as a representative species in naturally colonized animals [33]. Association of the murine ear skin with *M. pachydermatis* results in a pronounced induction of fungus specific Th17 cells in the skin and dLN [10]. This response was dependent on CD11c^+^ DCs, as we have previously shown [13]. The complex DC population of the skin comprises LCs in the epidermis as well as cDC1 and cDC2 subsets in the dermis [34]. To characterize DC dynamics and activation in response to *M. pachydermatis* skin colonization, we adapted a high parameter flow cytometry panel, established for murine skin [35]. The gating strategies for ear skin and dLN was consistent across the analyzed timepoints (**Fig. S3A-B).** Skin colonization with *M. pachydermatis* resulted in a massive infiltration of myeloid cells into the skin as early as one day post colonization, and further increased over the following two days (**Fig. 3A-C**). This infiltrate was composed primarily of neutrophils and monocytes (**Fig. 3C**). Skin DC numbers did not change markedly during the first three days of colonization, except in case of LCs, representing the largest of skin DC population prior to colonization, which declined slightly by day 2 after *M. pachydermatis* -association (**Fig. 3C**). Quantification of DC subsets in the skin-draining LN indicated that DCs readily migrated to the dLN on day two and three after fungal colonization, including cDC1, cDC2 and LCs (**Fig. 3A-B, D**). LN-resident cDC1s and cDC2s also slightly increased in numbers (**Fig. 3A-B, E**). Among the migratory DCs, cDC2s were the most prominent DC subset in the dLN (**Fig. 3D**), showing the strongest increase during colonization, followed by LCs while cDC1s only showed a minimal increase (**Fig. 3F**). Additionally, strong recruitment of macrophages, neutrophils, and monocytes to the dLN could be observed (**Fig. 3C-D**).

**Figure 3.**
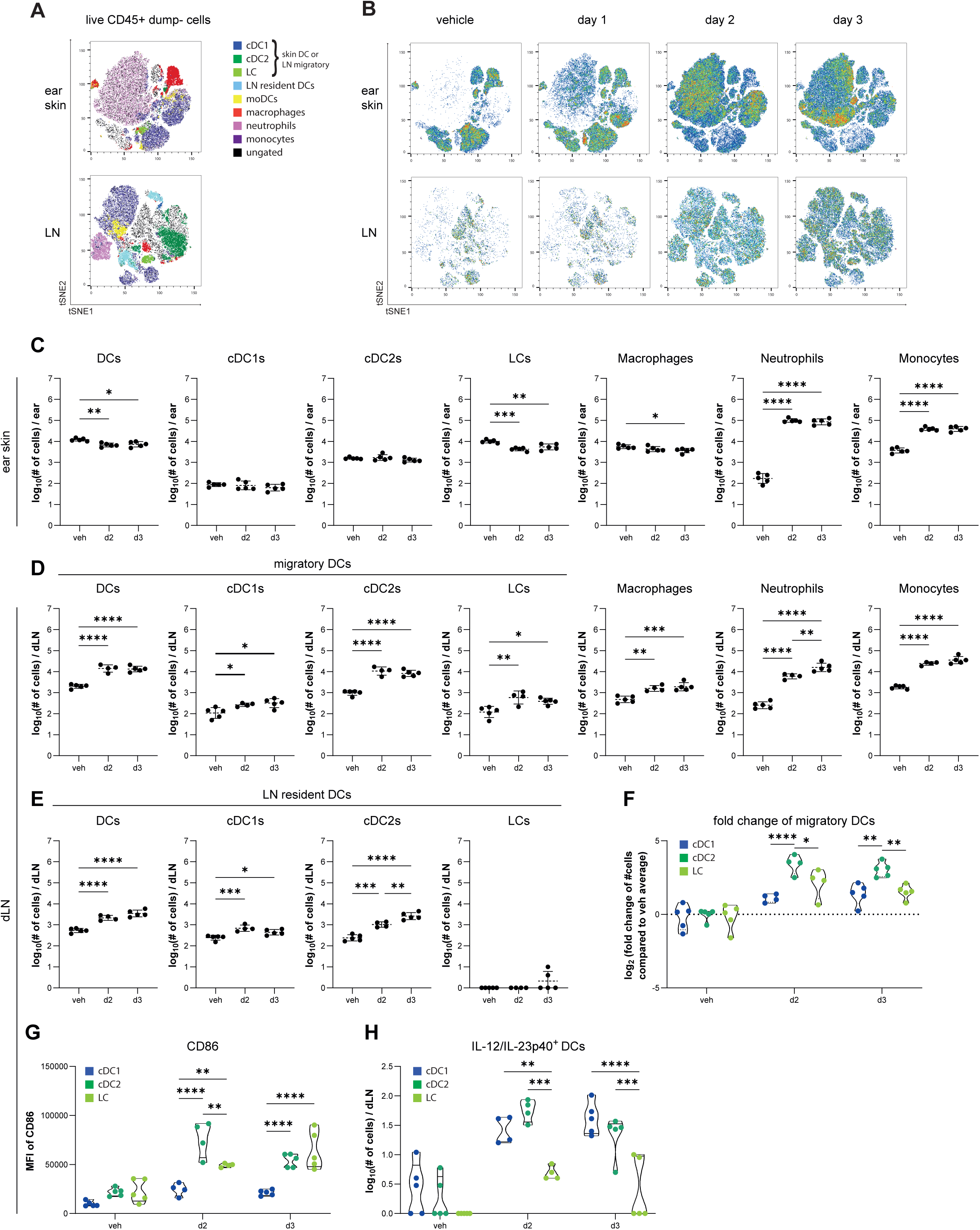
*Malassezia* skin colonization recruits myeloid cells and activates cDC2s. The ear skin of WT mice was colonized with *M. pachydermatis* for 1, 2 or 3 days as indicated or treated with vehicle (veh). **A.** tSNE analysis of myeloid cell populations in the ear skin and dLN of 15 concatenated samples including all conditions. **B.** tSNE analysis of all live CD45+ dump- cells in ear skin and dLN. The dump channel includes the markers CD3ε, NK1.1, and CD19. **C-E.** Quantification of myeloid cells in the ear skin (**C**) and dLN (**D-E**). **F.** Fold change of migratory DC subsets in the LN. Log2 fold changes were calculated using the mean of the vehicle control for each DC subset. **G.** MFI of CD86 in migratory DC subsets in the dLN. **H.** Quantification of IL-12/IL-23p40^+^ cells among migratory DC subsets in the dLN. n= 5, 4, 5, data from one representative of three independent experiments, mean ± SD. Statistical significance was determined using one-way ANOVA, * p < 0.05, ** p < 0.01, *** p<0.001, **** <0.0001. **See also Fig. S3 and Fig. S4.**

To obtain further insights into the DC response to *Malassezia*, we next examined the activation state of the migratory DC subsets in the dLN. cDC2s exhibited most pronounced elevation in CD86 expression at day two and day three after *M. pachydermatis* colonization (**Fig. 3G-H, Fig. S3C**). LCs also exhibited high CD86 expression, albeit with slower kinetics (**Fig. 3G**). Likewise, when looking at cytokine production, cDC2s were the most prominent source of IL-12/23p40 at day two after colonization (**Fig. 3H, Fig. S3D**). Consistently, Batf3- dependent cDC1s including Langerin^+^ DCs and LCs were redundant for Th17 induction in response to fungal skin colonization (**Fig. S4A-F**). Together, these data imply relevance for cDC2s in the cutaneous response to *Malassezia*, and in the activation of protective type 17 immunity in particular.

### Dectin-2, but not Mincle nor Dectin-1, mediates Th17 immunity to *Malassezia*

Diverse lymphocyte subsets contribute to protective IL-17 immunity against *Malassezia* [10], [13]. While dermal γδ T cells represent a major source of IL-17 from the first days of colonization, αβ T cells, and CD4^+^ αβ T cells in particular, also contribute significantly to the overall IL-17 response, especially from 7 days after colonization [13]. The relevance of αβ T cells is emphasized when comparing fungal control in mice lacking γδ T cells (*Tcrd*^-/-^) and mice lacking both γδ T and αβ T cells (*Tcrbd*^-/-^). While overall numbers of CD90^+^ cells producing IL-17A in response to *M. pachydermatis* were not, or only partially, affected by the lack of γδ T cells in *Tcrd*^-/-^ mice in comparison to WT controls, the additional lack of αβ T cells in *TCRbd*^-/-^ mice reduced the IL-17A response to nearly baseline levels in the skin and dLN on day 12 after colonization (**Fig. 4A-B**). The relevance of αβ T cells in the anti- *Malassezia* response was further highlighted by the increased skin fungal burden in *Tcrbd*^-/-^ compared to *Tcrd*^-/-^ mice (**Fig. 4A, right**) and the increased numbers of IL-17 producing CD4^+^ T cells in the ear (**Fig. 4A, middle**), suggesting that these cells compensate for the lack of γδ T cells in *Tcrbd*^-/-^ mice for producing IL-17. The relevance of αβ T cells in antifungal immunity is reminiscent of what has been described in humans where *Malassezia*-responsive memory CD4^+^ T cells belong primarily to the Th17 subset [10], [11].

**Figure 4.**
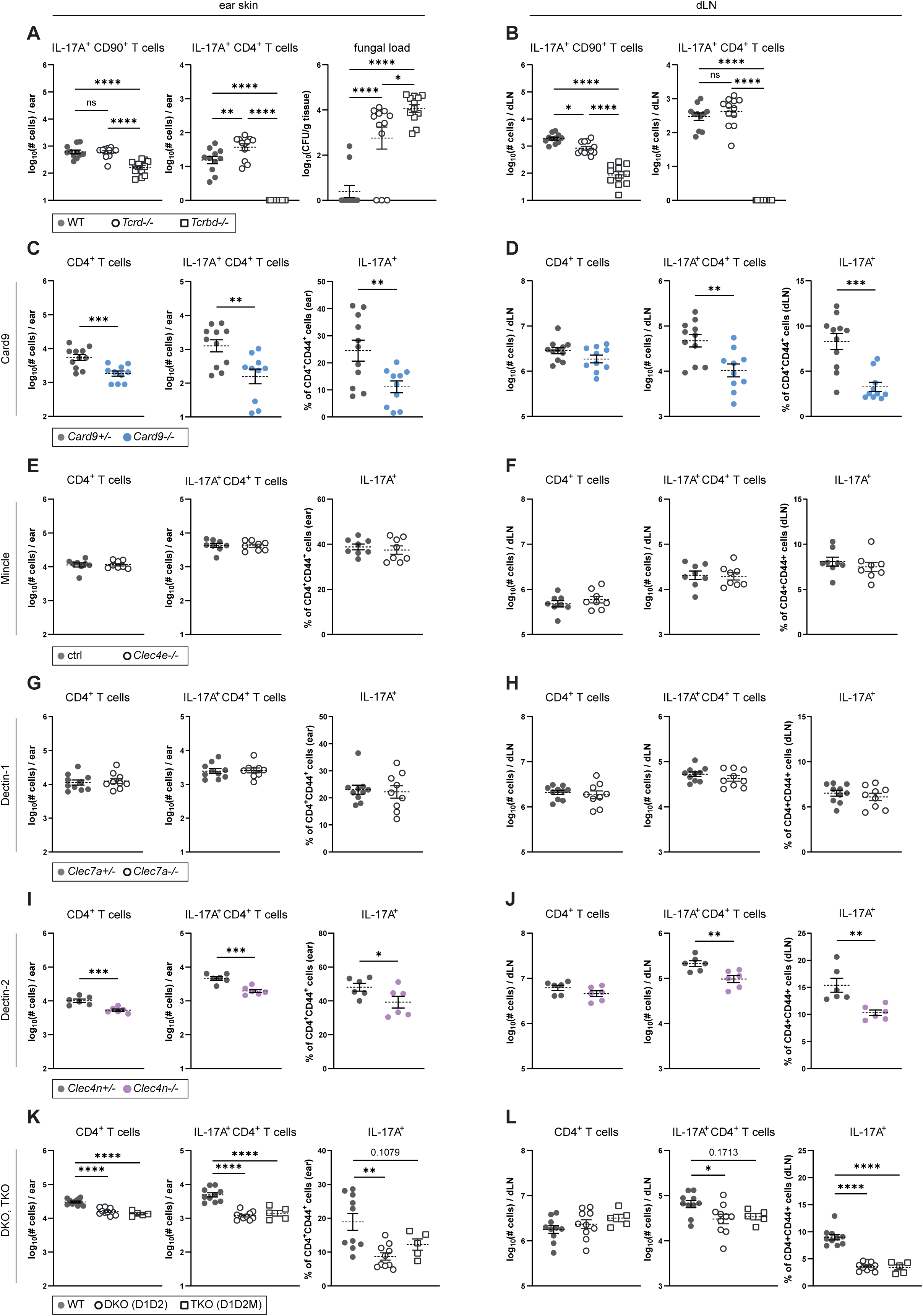
Dectin-2 but not Mincle nor Dectin-1 mediates Th17 immunity to *Malassezia*. **A-B.** The ear skin of *Tcrd*^-/-^, *Tcrbd*^-/-^ and WT mice was colonized with *M. pachydermatis* for 12 days. IL-17A^+^ CD90^+^ cell counts and IL-17A^+^ CD4^+^ cell counts were quantified in ear skin (A) and dLN (B). Fungal load (CFU) was assessed in ear skin. n = 11, 12, 12, data pooled from three independent experiments, mean ± SEM. **C-L.** The ear skin of *Card9*^-/-^ and *Card9*^+/-^ mice (C, D), *Clec4e*^-/-^ and control chimeras (E, F), *Clec7a*^-/-^ and *Clec7a*^+/-^ mice (G, H), *Clec4n*^-/-^ and *Clec4n*^+/-^ mice (I, J) and Dectin-1-Dectin-2 (DKO) and Mincle-Dectin-2-Dectin-1 (TKO) chimeras (K, L) was colonized with *M. pachydermatis* for 7 days. CD4^+^ T cell counts, IL-17A^+^ CD4^+^ CD44^+^ T cell counts, and the % of IL-17A producing CD4^+^ CD44^+^ T cells were quantified in the ear skin (C, E, G, I, K) and dLN (D, F, H, J, L). C, D: n = 11, 10, data pooled from three independent experiments, mean ± SEM; E, F: n = 8, 8, data pooled from three independent experiments, mean ± SEM; G, H: n = 10, 9, data pooled from two independent experiments, mean ± SEM; I, J: n = 6, 6, female mice only, data pooled from two independent experiments, mean ± SEM; K, L: n = 10, 10, 5, pooled from two independent experiments, mean ± SEM. Statistical significance was determined using one-way ANOVA (A-B, K-L) or unpaired t test (C-J) or, * p < 0.05, ** p < 0.01, *** p<0.001, **** <0.0001. **See also Fig. S5 and Fig. S6**.

To investigate the role of the CLR pathway in the regulation of the Th17 response to *Malassezia*, we first assessed Th17 induction in dependence on Card9. For this, we colonized *Card9*^-/-^ and heterozygous littermate control mice with *M. pachydermatis* and analyzed the αβ T cell response in ear skin and dLN 7 days later. To quantify cytokine production, CD4^+^ T cells from skin and dLN were re-stimulated with PMA and ionomycin or with heat-killed fungus pulsed DCs, respectively, and analyzed by flow cytometry (**Fig. S5A**). CD4^+^ T cells and the IL-17A-producing subset of activated CD4^+^ CD44^+^ cells in particular were reduced in skin and dLN of *Card9*^-/-^ compared to littermate control mice (**Fig. 4C-D**). This was true when quantifying cell numbers as well as percentages of IL-17A producing CD4^+^ CD44^+^ T cells (**Fig. 4C-D**). IL-22 producing CD4^+^ T cells and IL-17A-IL-22 double producers were also reduced in absence of Card9 (**Fig. S5B-C**). The observed Card9-dependence of the Th17 response was specific to *Malassezia* skin colonization and not a consequence of dysregulated immune homeostasis, as no baseline differences in overall and IL-17A-producing CD4^+^ T cells in skin and dLN could be revealed between non-colonized *Card9*^-/-^ and *Card9*^+/-^ animals (**Fig. S5D-E**). We then turned towards interrogating which of the CLRs mediating DC activation in response to *Malassezia* (**Fig. 2C-E**) was involved in coupling innate fungal recognition to Th17 induction. To study the role of Mincle we generated radiation chimeras in which the hematopoietic compartment was restored with bone marrow from *Clec4e*^-/-^ (or WT control) mice as live *Clec4e*^-/-^ mice were not available to us (**Fig. S5F**). The Th17 cell response to *M. pachydermatis* was not altered in skin and dLN of *Clec4e*^-/-^ chimeras and overall CD4^+^ T cells as well as the IL-17A- and IL-22 producing subsets remained unchanged when compared to control chimeras (**Fig. 4E-F, Fig. S5G-H**). The same was observed in *M. pachydermatis* - colonized *Clec7a*^-/-^ mice in comparison to their heterozygous littermates as controls (**Fig. 4G-H, Fig. S5I-J**). These results were surprising considering the pronounced requirement of both Mincle and Dectin-1 for full DC activation *in vitro* and this led us to speculate about a general redundancy of individual CLRs for *Malassezia* immunity *in vivo*. Analysis of *Clec4n*^-/-^ mice however disproved this hypothesis. Overall CD4^+^ T cells as well as the IL-17A- and IL-22 producing populations in particular, were reduced in skin and dLN of *Clec4n*^-/-^ mice when compared to littermate controls (**Fig. 4I-J, Fig. S6A-B**) revealing a non-redundant role of Dectin-2 in the Th17 response to *Malassezia in vivo*, although the effect appeared more prominent in female than male animals (**Fig. S6C-D**). The relevance of Dectin-2 for Th17 induction in *M. pachydermatis-* colonized skin and dLN was confirmed in another species of *Malassezia*, *M. sympodialis* (**Fig. S6E-F**), while Dectin-1 remained redundant in response to this species as well (**Fig. S6G-H**). Comparable Th17 responses in *Clec4n*^-/-^ and *Clec4n*^+/-^ mice at steady state confirmed that the observed Dectin-2 dependence of *Malassezia*-responsive Th17 immunity was not a consequence of baseline differences between the two groups (**Fig. S6I-J**).

We speculated that a putative contribution of Dectin-1 or Mincle to the Th17 response to *Malassezia* might be masked in the presence of fully functional Dectin-2. We therefore tested the consequences of concomitant lack of two or three CLRs. For this, we generated Dectin-1-Dectin-2 DKO and Mincle-Dectin-2-Dectin-1 TKO chimeric mice [32] (**Fig. S6K**). When colonizing these chimeras with *M. pachydermatis*, we reproduced a pronounced reduction of the Th17 response in both DKO and TKO chimeric mice (**Fig. 4K-L**). To assess the level of IL-17 reduction we compared the fold changes of average numbers of IL-17A^+^ CD4^+^ T cells between knockout and control mice. The reduction of Th17 cells in DKO and TKO mice was comparable (ear skin: DKO: 0.23, TKO: 0.28 – dLN: DKO: 0.52, TKO: 0.48) confirming the redundant role of Mincle observed in the *Clec4e*^-/-^ mice. Comparing the fold reduction of IL-17A^+^ CD4^+^ T cell numbers between Dectin-2 deficient animals (ear skin: *Clec4n*^-/-^: 0.42 – dLN: *Clec4n*^-/-^: 0.47) and DKO and TKO mice revealed a partial contribution of Dectin-1 especially in the skin in addition to Dectin-2. *Card9*^-/-^ mice showed a stronger reduction of IL-17A^+^ CD4^+^ T cells in the ear skin and dLN than any of the CLR knockout mice (ear skin: *Card9*^-/-^: 0.14, – dLN: *Card9*^-/-^: 0.23). Together, these data identify Dectin-2 as the most prominent Card9- dependent CLR for activating protective Th17 immunity in response to *Malassezia* spp. recognition.

### The Th17 response against *Malassezia* depends on T-cell-intrinsic MyD88 signaling

Complementary to CLRs, we also considered the contribution of other PRR families in the activation of adaptive immunity against *Malassezia*. Our RNA-Seq dataset revealed upregulation of multiple Toll-like receptors (TLRs) such as TLR2, TLR7, TLR8, and TLR9 in the colonized skin (**Fig. 1A**). TLRs have been implicated in fungal recognition including the recognition of *Malassezia* [36], [37]. We first assessed the impact of TLR and MyD88 deficiency on *Malassezia* -induced DC activation. Stimulation of BMDCs generated from *Tlr23479*^-/-^ mice with *M. pachydermatis*, *M. sympodialis* and *M. furfur* resulted in reduced cytokine secretion compared to WT controls (**Fig. S7A**). Consistent with this result, MyD88- deficient BMDCs were strongly impaired in cytokine production in response to *Malassezia* spp. (**Fig. S7B**) while TLR and MyD88 deficiency was confirmed with CpG and both cells responded to curdlan (β-1,3 glucan) when compared to the unstimulated control, as expected (**Fig. S7A-B**). To assess the relevance of TLR- and MyD88-dependent signaling for *Malassezia*-induced Th17 immunity, we quantified the CD4^+^ T cell response in ear skin and dLN of *M. pachydermatis* colonized *Tlr23479*^-/-^ and *Myd88*^-/-^ deficient mice. While TLR deficiency had no impact on the antifungal Th17 response (**Fig. 5A-B**), overall CD4^+^ T cells and IL-17A and IL-22 producing CD4^+^CD44^+^ T cells were strongly reduced in *MyD88*^-/-^ mice when compared to heterozygous littermate controls (**Fig. 5C-D, Fig. S7C-D**). Th17 responses at steady state were assessed to exclude putative baseline differences in homeostatic immunity due to constitutive deletion of *Myd88*, whereby no differences in overall or IL-17A producing CD4^+^ T cells could be detected between non-colonized *MyD88*^-/-^ animals and littermate controls (**Fig. S7E-F**). Together, these results suggest a role for IL-1 family cytokines, which also signal via MyD88 [38], in the cutaneous Th17 response to *Malassezia*. In contrast to C-type lectin receptors, whose expression is largely restricted to myeloid cells, MyD88 is widely expressed in hematopoietic and non-hematopoietic cells. To decipher in which cellular compartment MyD88 is required to mount a protective Th17 response to *Malassezia*, we generated bone marrow chimeric mice by reconstituting irradiated CD45.2^+^ MyD88-deficient and CD45.1^+^ WT mice with CD45.1^+^ WT or CD45.2^+^ MyD88-deficient bone marrow cells, respectively (**Fig. S7G**). Overall CD4^+^ T cell numbers were comparable in skin and dLN of all experimental groups after colonization with *M. pachydermatis* (**Fig. 5E-F**). MyD88 was essential in the hematopoietic compartment for optimal IL-17A production by CD4^+^ T cells, while the contribution in the radioresistant compartment was more variable and varied between skin and dLN (**Fig. 5E-F**).

**Figure 5.**
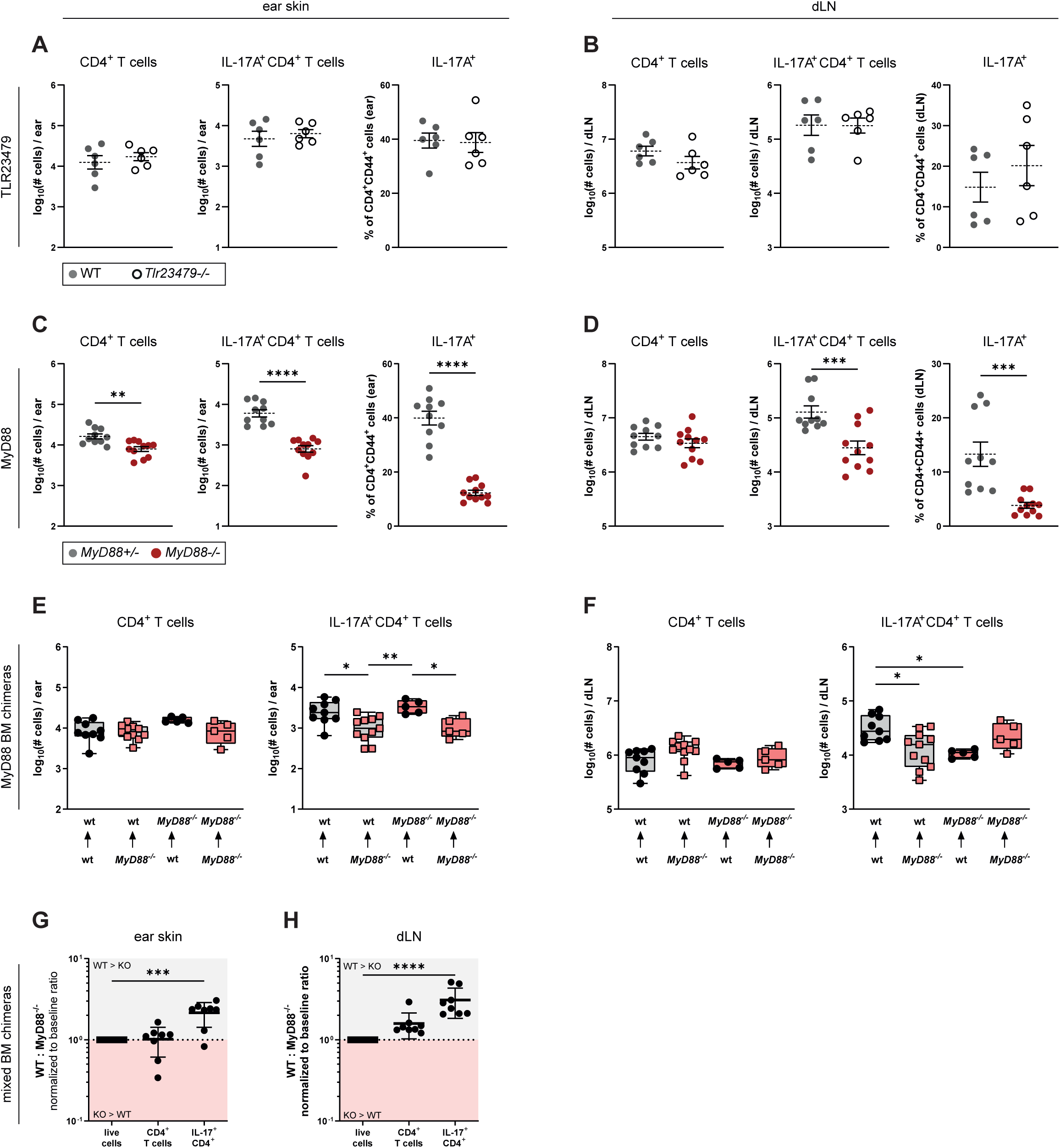
The Th17 response against *Malassezia* depends on T-cell-intrinsic MyD88 signaling. **A-D.** The ear skin of *Tlr23479*^-/-^ and WT mice (A, B) and *MyD88*^-/-^ and *MyD88*^+/-^ mice was colonized with *M. pachydermatis* for 7 days. CD4^+^ T cell counts, IL-17A^+^ CD4^+^ CD44^+^ T cell counts, and the % of IL-17A producing CD4^+^ CD44^+^ T cells were quantified in ear skin (A, C) and dLN (B, D) A, B: n = 6, 6, data pooled from two independent experiments, mean ± SEM; C, D: n = 10, 11, data pooled from three independent experiments, mean ± SEM. **E-F.** Chimeric mice were generated by irradiating WT (grey box) or *MyD88*^-/-^ (red box) hosts that were reconstituted with WT (black circles) or *MyD88*^-/-^ (red squares) bone marrow. CD4^+^ T cell counts, IL-17A^+^ CD4^+^ CD44^+^ T cell counts, and the % of IL-17A producing CD4^+^ CD44^+^ T cells were quantified in ear skin (E) and dLN (F). n = 9, 11, 5, 5, data pooled from three independent experiments, mean ± SEM. **G-H.** Mixed BM chimeras were generated by reconstituting WT hosts with a 1:1 mix of WT and *MyD88*^-/-^ bone marrow. The ratio of WT to *MyD88*^-/-^ CD4^+^ T cells and IL-17A producing CD4^+^ T cells in the ear skin (G) and dLN (H) was calculated by normalization to the baseline ratio of WT to *MyD88*^-/-^ cells. n = 8,8, data pooled from two independent experiments, mean ± SEM. Statistical significance was determined using unpaired t test (A-D) or one-way ANOVA (**E-H**), * p < 0.05, ** p < 0.01, *** p<0.001, **** <0.0001. **See also Fig. S7.**

Based on these observations, we hypothesized that MyD88 may act in a T cell-intrinsic manner. To test this, we generated mixed bone marrow chimeras. WT mice were irradiated and reconstituted with a 1:1 mixture of CD45.2^+^ MyD88-deficient and CD45.1^+^ WT bone marrow cells (**Fig. S7H**). While the ratio of WT to *Myd88*^-/-^ CD45^+^ cells overall remained stable during colonization (**Fig. S7H**), the ratio within the total CD4^+^ T cell population, and the IL-17A^+^ CD4^+^ T cell population was skewed towards the WT genotype in both skin and dLN (**Fig. 5G-H**). Together, these data indicate that *Malassezia*-specific Th17 responses depend on MyD88 in the hematopoietic compartment and further implicate a T-cell intrinsic requirement of MyD88-dependent signaling in this context. Furthermore, the redundancy of TLRs for Th17 immune induction suggests that IL-1 family cytokines are involved.

### *Malassezia-*induced inflammatory IL-1 signaling induces Th17 responses

Besides its role in the TLR pathway, MyD88 also mediates IL-1 family cytokine signaling. Scrutinizing again our RNA-Seq dataset revealed significant upregulation of numerous IL-1 family cytokines in the *M. pachydermatis* -colonized murine skin (**Fig. 6A**). Among the top candidates were *Il1b* (encoding IL-1β), *Il1f6* (encoding IL-36α), *Il1f9* (encoding IL-36γ), *Il1f8* (encoding IL-36β) and *Il1a* (encoding IL-1α) (**Fig. 6A**). Consistent with the inflammatory nature of these cytokines, we observed a MyD88-dependent increase in inflammation in the colonized ear in histological examinations. Cutaneous hyperplasia peaked 4 days after colonization, which was reduced again by day 7 after colonization (**Fig. 6B**), as we showed previously [13].

**Figure 6.**
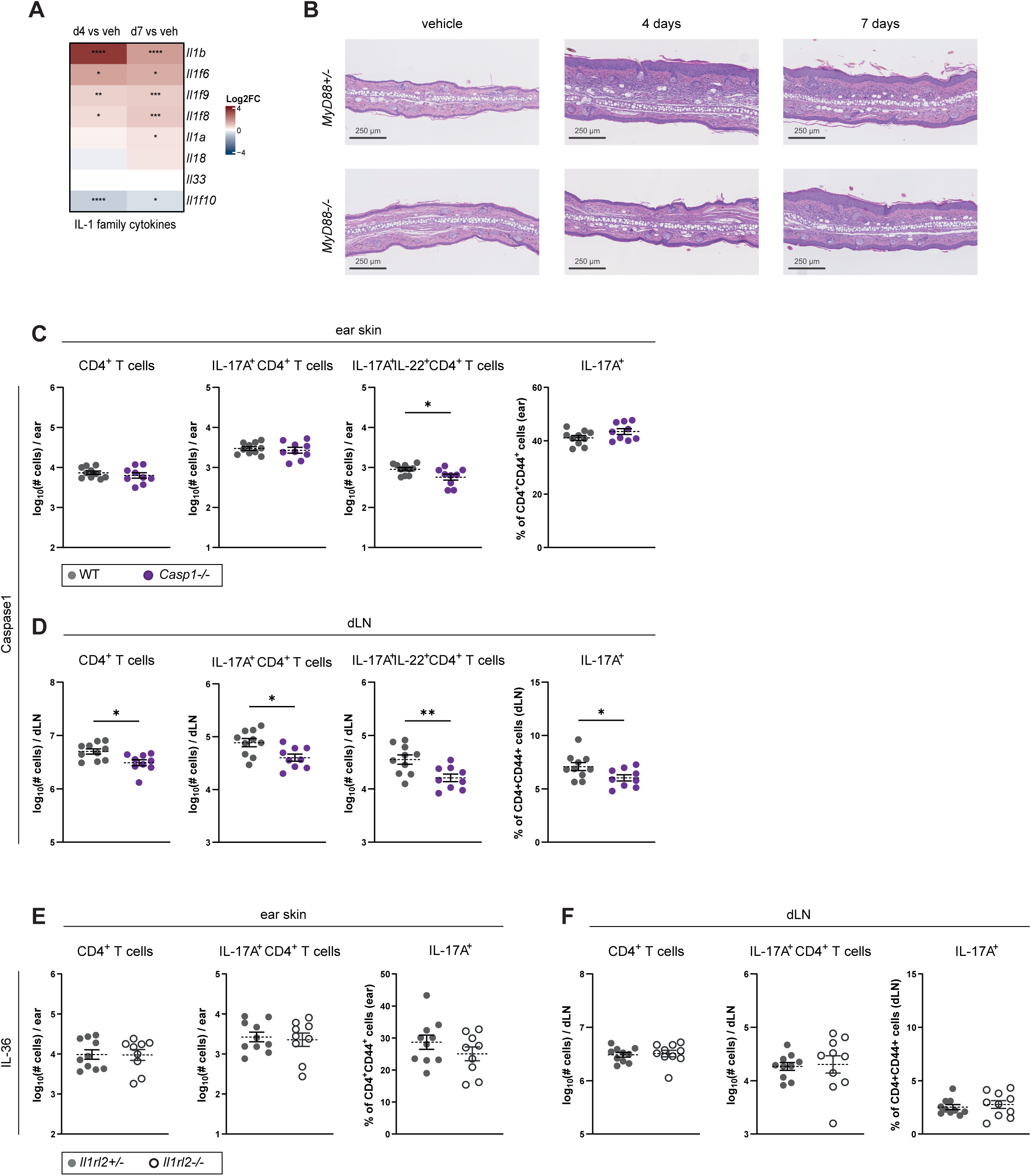
*Malassezia*-induced inflammatory IL-1 signaling induces Th17 responses. **A.** Differentially regulated genes (log2 fold change) of the IL-1 family cytokine genes in the ear skin of mice that have been colonized for 4 days (d4) or 7 days (d7) with *M. pachydermatis* or treated with vehicle (veh). **B.** Hematoxylin and eosin-stained ear sections from *MyD88*^-/-^ and *MyD88*^+/-^ mice after 4 or 7 days of colonization with *M. pachydermatis*. **C-F.** The ear skin of *Caspase1*^-/-^ and WT mice (C, D) and *Il1rl2*^-/-^ and *Il1rl2*^+/-^ mice (E, F) was colonized with *M. pachydermatis* for 7 days. CD4^+^ T cell counts, IL- 17A^+^ (and IL-17A^+^ IL-22^+^) CD4^+^ CD44^+^ T cell counts, and the % of IL-17A producing CD4^+^ CD44^+^ T cells were quantified in ear skin (C) and dLN (D). C, D: n = 10, 10, data pooled from two independent experiments, mean ± SEM); E, F: n = 10, 10, data pooled from two independent experiments, mean ± SEM. Statistical significance was determined using unpaired t test, * p < 0.05, ** p < 0.01. **See also Fig. S8**.

To test whether IL-1 family cytokine signaling via MyD88 was required for the induction of Th17 immunity in response to *Malassezia*, we made use of caspase-1 deficient mice. Caspase-1 cleaves inactive pro-IL-1β to release bioactive IL-1β. It is also required for the maturation of IL-18 and IL-37 [39] and at least partially for IL-1α production [40]. The T cell cytokine production in response to *M. pachydermatis* was indeed reduced in skin and dLN of *Caspase1*^-/-^ mice (**Fig. 6C-D, Fig. S8A-B**) confirming the relevance of IL-1 family cytokines in the induction of the cutaneous Th17 response to the fungus. In contrast and opposed to a previous report[41], IL-36 receptor deficiency did not impair IL-17 responses in skin and dLN (**Fig. 6E-F, Fig. S8C-D**). Together, these data support that cutaneous Th17 immunity elicited by *Malassezia* depends on a T-cell-intrinsic, caspase-1-dependent, but IL-36-independent signal that complements Dectin-2/Card9-induced cues.

## DISCUSSION

Th17 cells orchestrate barrier immunity to maintain tissue homeostasis and stable microbial colonization. As such, Th17 cells also contribute to the immunosurveillance of *Malassezia* spp., which make up by far the largest and most widespread members of the fungal community of the skin microbiome. Induction of Th17 immunity relies on PRR activation in DCs that couple microbial recognition to T cell priming for the generation of host-protective effector T cells. Here, we show that *Malassezia* is recognized by the three distinct CLRs – Mincle, Dectin-1, and Dectin-2, whereby only the latter is required *in vivo* for the induction of *Malassezia*-specific IL-17A producing CD4^+^ T cells. This exceeding role of Dectin-2 and the redundancy of Mincle was confirmed by the comparison of Dectin-2^-/-^ with Dectin-1-Dectin-2 DKO and Mincle-Dectin-2-Dectin-1 TKO mice.

*O*-linked mannoprotein was proposed to serve as a ligand for Dectin-2 in *Malassezia* [42], although its exact molecular identity remains to be determined. Our observation that the antifungal response is Dectin-2 dependent for at least two different species of *Malassezia* suggests that the ligand may be conserved within the genus. Mannan is a common Dectin-2 agonist in other fungi and non-fungal microbes that are recognized by this CLR [43], [44], [45].

The most well-known ligand of Dectin-1 is β-1,3-glucan [46], though some other ligands, such as annexin and N-glycan, have also been described [47]. In the cell wall of *Malassezia,* β-1,6-glucan is far more abundant than β-1,3-glucan [48]. The relevance thereof in the context of *Malassezia* skin colonization and with respect to Dectin-1 engagement remains to be determined, although Dectin-1 has been reported to be able to recognize β-1,6-glucan [49].

The redundant role of Mincle in Th17 immunity against *Malassezia* was somewhat surprising considering previous studies that identified Mincle as a potent immune receptor for *Malassezia* [21], [42], [50] and attributed Mincle an important role in *Malassezia* phagocytosis by macrophages [51]. While we confirmed these results with isolated DCs in culture, the situation is different *in vivo*. BMDCs only partially reflect tissue-resident DCs in the skin where a more complex network of immune signaling pathways may compensate for the lack of individual CLRs. We cannot fully exclude a role of Mincle in LCs, which are radioresistant and remain of host origin in radiation chimeras [52], given that we used chimeras for the study of Mincle *in vivo*. However, this is unlikely since LCs were not found to be essential for the induction of the Th17 response against *Malassezia*.

The cell wall is a moving target for the immune system and its composition and the exposure of specific PAMPs underlies environmental cues, such as nutrient availability. As such, changes in carbon sources, hypoxia or iron depletion modulate the degree of β-glucan exposure in *Candida albicans* via a mechanism termed “β-glucan shaving” [53], [54], [55], [56]. Morphological changes, such as the yeast-to-hyphae transition in *C. albicans* [57] or the germination of *A. fumigatus* conidia [58], [59] also significantly impact the cell wall composition and structure. By these mechanisms, fungi change the availability of specific CLR ligands, and thereby create the need for a diverse CLR recognition network in the host, which may appear redundant if analyzing them under a single condition. Whether the cell wall of *Malassezia* also displays plasticity under different environmental conditions has not been addressed. It is tempting to speculate that when exposed to different skin conditions such as in atopic dermatitis where lipid composition and pH differ largely from healthy skin [60], [61], [62], [63], *Malassezia* undergoes changes in its cell wall and thus in CLR recognition, which impact the quality of the elicited immune response. Such mechanisms may explain the commonly observed sensitization against *Malassezia* in atopic dermatitis patients. In fact, *Malassezia*-specific IgE antibody levels correlate with disease severity, and this is most pronounced in the head-and-neck subtype of disease [64], while fungus-reactive Th2 cells are thought to contribute to pathogenesis via a mechanism involving cross-reactivity to host proteins [65], [66].

Our study further identified MyD88-dependent signaling as a requirement for Th17 immunity against *Malassezia*. We attributed this at least in part to Caspase-1 dependent IL-1 family cytokines while excluding IL-36 cytokines and TLR2, TLR3, TLR4, TLR7 and TLR9. The redundancy of TLRs for Th17 induction by *Malassezia* was not mirrored in vitro, where TLRs were also required for maximal DC activation. The involvement of IL-1 family cytokines is not unexpected considering that *Malassezia* colonization of murine ear skin induces mild inflammation [13]. Caspase-1 dependent IL-1 family cytokines include IL-1β, IL-33, IL-18, and IL-37, whereby there is no homologue of IL-37 in mice [67], and IL-1α secretion depends at least partially on Caspase-1 [68]. IL-1 has been implicated in Th17 differentiation [69] by its capacity to downregulate FoxP3 [70], [71]. IL-1 is also involved in eliciting IL-17 production by innate source of IL-17, including dermal γδ T cells, as we recently showed in the context of *Malassezia*-colonized skin [13]. Our finding that IL-36 family cytokines are redundant for antifungal Th17 immunity in the skin contrasts with a previous report [41]. The reason for the discrepancy remains unclear but may be linked to differences in the experimental model, the fungal species studied, the time point analyzed, and/or the degree of detail with which the type 17 response was analyzed. Additionally, an effect of haploinsufficiency in the Il1rl2^-/-^ mice cannot be fully excluded (oral communication, M. Kopf). Independence of the IL-17 response from IL-36 has also been reported for other fungal barrier infection models [72]. An involvement of other MyD88-dependent IL-1 family receptors such as Il1rl1 in *Malassezia*-induced Th17 induction cannot be excluded. IL-1 family cytokines exert functions beyond the regulation of type 17 responses, which may be of particular relevance in the context of *Malassezia*-associated inflammatory skin diseases such as atopic or seborrheic dermatitis, as supported by the MyD88-dependence of the *Malassezia*-induced epidermal hyperplasia and with potential implications for therapy [73].

Complementary to αβ T cells, γδ T cells also contribute to the overall IL-17 response to *Malassezia*, although with different kinetics [10], [13]. Surprisingly, γδ T cells are activated independently of Card9 and CLR signaling [13]. Instead, they were activated by IL-23 and IL-1 cytokines and respond to soluble *Malassezia* metabolites once licensed [13]. By following distinct modes of activation, αβ T cells and γδ T cells complement each other thereby emphasizing the robustness of the response in line with the relevance of the type 17 response for fungal control [74], [75], [76], [77]. The contribution of the Th17 vs. γδ T17 cells may be of particular interest in humans as γδ T cells in human skin are inferior to those in murine skin regarding their IL-17 producing capacity [78]. To what extent CD8^+^ T cells, which also produce IL-17 in the murine skin [13], contribute to *Malassezia* control and how their effector functions are activated and regulated remains to be determined.

Taken together, our work demonstrates that during skin colonization with the abundant fungal commensal *Malassezia*, a complex array of immune pathways and cellular players elicits a robust response to control fungal colonization, preventing overgrowth and ensuring skin homeostasis. By acting in two distinct cellular compartments, Card9 and MyD88 act as central coordinators of the Th17-mediated immunosurveillance mechanism. Elucidating the role and relevance of these pathways in the antifungal response under inflamed skin conditions will help understanding the mechanism of pathogenesis in atopic dermatitis and other prevalent chronic-inflammatory skin conditions that are associated with *Malassezia* colonization.

## DATA LIMITATIONS AND PERSPECTIVES

While the experimental model of *Malassezia* skin colonization bears great potential to study *Malassezia*-host interactions *in vivo*, in the context of a functional immune system including tissue-resident and infiltrating cells, the murine skin exhibits important differences to human skin. Moreover, the primary exposure of mice kept under specific pathogen free conditions to *Malassezia* elicits an inflammatory response characterized by infiltration of inflammatory cells, unlike the situation during commensal colonization of healthy human or animal skin. In turn the fungus does not persist on murine skin but is cleared after 2 weeks in wild type mice [10]. We conducted most *in vivo* experiments with a single species of *Malassezia* although key findings were confirmed with a second species. Whether our findings can be generalized for the entire genus of *Malassezia*, and for the most abundant human colonizing species *M. restricta* and *M. globosa* in particular, remains to be demonstrated.

## MATERIALS AND METHODS

### Ethics approval statement for animal studies

All mouse experiments in this study were conducted in strict accordance with the guidelines of the Swiss Animals Protection Law and were performed under the protocols approved by the Veterinary office of the Canton Zurich, Switzerland (license number 168/2018 and 142/2021). All efforts were made to minimize suffering and ensure the highest ethical and humane standards according to the 3R principles [101].

### Animals

WT C57BL/6j mice were purchased from Janvier Elevage. Ly5.1 [79], *TCRd*^-/-^ [80] *TCRbd*^-/-^ [80], [81], *Card9*^-/-^ [10], *MyD88*^-/-^ [82], *Clec4n*^-/-^ [83] and *Clec7a*^-/-^ [84] (kindly provided by Gordon Brown, University of Exeter, UK), *Tlr23479*^-/-^ [85] (kindly provided by Thorsten Buch, University of Zürich, Switzerland), *Il1rl2*^-/-^ (kindly provided by Manfred Kopf, ETH Zürich, Switzerland), *Batf3*^-/-^ mice [86] (kindly provided by Mark Suter, University of Zurich, Switzerland and Bavarian Nordic), and Langerin-DTR mice [87] (kindly provided by Marc Vocanson, Inserm, Lyon, France), were bred at the Institute of Laboratory Animals Science (LASC, University of Zürich, Switzerland). *Casp1*^-/-^ [88] and associated WT control mice (kindly provided by Wolf-Dietrich Hardt, ETH Zürich, Switzerland) were bred at the ETH Phenomics Center (EPIC, ETH Zürich, Switzerland). All mice were on the C57BL/6 background. The animals were kept in specific pathogen-free conditions and used at 8-14 weeks of age in age-matched groups. Female and male mice were used for experiments unless otherwise specified.

### DTR treatment

Homozygous Langerin-DTR mice were injected i.p. with 1 µg diphtheria toxin (Sigma-Aldrich/Merck) or PBS control one day prior to *Malassezia* colonization and on the day of colonization as previously described [89], [90]. Depletion of LCs was assessed in ears and dLN of colonized mice after 7 days.

### Generation ofh chimeric mice

C57BL/6j WT, Ly5.1 or *MyD88*^-/-^ female recipient mice at 6-8 weeks of age were irradiated twice with a dose of 5.5 Gy at an interval of 12 h. *Clec4e*^-/-^ mice ([91]kindly provided by David Sancho, CNIC, Spain), Dectin-1-Dectin-2 DKO [32], Mincle- Dectin-2- Dectin-1 TKO mice [32] or *MyD88*^-/-^ and WT C57BL/6 or Ly5.1 controls respectively, served as bone marrow donors. For mixed BM chimeras, recipient mice were reconstituted with a 1:1 mix of C57Bl/6 and *MyD88*^-/-^ bone marrow. The bone marrow of one donor mouse was injected in the tail vein of five recipient mice each, 6 h after the second irradiation. Mice were treated with Borgal® (MSD Animal Health GmbH) p.o. for the first 2 weeks of an 8- week reconstitution period.

### Fungal strains

**A.** *M. pachydermatis* strain ATCC 14522 [92] (CBS 1879), *M. sympodialis* strain ATCC 42132 [93] and *M. furfur* strain JPLK23 [92] (CBS 14141) were grown in mDixon medium at 30°C and 180 rpm for 2-3 days. Heat-killing was achieved by incubating fungal suspensions at a concentration of 5×10^6^ CFU/ml in PBS for 45 min at 85°C.

### Epicutaneous colonization of mice with *Malassezia*

Epicutaneous colonization of the mouse ear skin was performed as described previously [10]. In short, *Malassezia* cells were washed with PBS and suspended in commercially available native olive oil. A 100 µl suspension containing 1x 10^7^ yeast cells was topically applied onto the dorsal skin of each ear while mice were anaesthetized. Animals treated with olive oil (vehicle) and infected animals were kept separately to avoid fungal transmission.

### Isolation of skin and lymph node cells

For digestion of total ear skin, mouse ears were cut into small pieces and transferred into Ca^2+^- and Mg^2+^-free Hank’s medium (Life Technologies) supplemented with Liberase TM (0.15 mg/mL, Roche) and DNAse I (0.12 mg/mL, Sigma-Aldrich) and incubated for 50 min at 37°C. Ear draining lymph nodes (dLN) were digested with DNAse I (2.4 mg/ml Sigma-Aldrich) and Collagenase I (2.4 mg/ml, Roche) in PBS for 30 min at 37°C. Both cell suspensions were filtered through a 70 μm cell strainer (Falcon) and rinsed with PBS supplemented with 5 mM EDTA (Life Technologies) and 1 % fetal calf serum.

### Ex vivo T cell re-stimulation

For *in vitro* re-stimulation of T cells, skin cell suspensions were incubated in a U-bottom 96-well plate (cells from 1/6 ear per well) with cRPMI medium (RPMI with L-Glutamine, Gibco) supplemented with fetal calf serum (10%, Omnilab, HEPES (10 mM, Gibco), sodium pyruvate (1X, Gibco), non-essential amino acids (1X, Gibco), β- mercaptoethanol (50 μM, Gibco), Penicillin (1%) and Streptomycin (1%) with phorbol 12- myristate 13-acetate (PMA, 50 ng/ml, Sigma-Aldrich) and ionomycin (500 ng/ml, Sigma-Aldrich) for 5 h at 37 °C in the presence of Brefeldin A (10 μg/ml). 1×10^6^ lymph node cells per well of a flat-bottom 96-well plate were co-cultured with 1×10^5^ DC1940 cells [94] that were previously pulsed with 2.5×10^5^ heat-killed fungal cells for 2 h. Brefeldin A (10 mg/ml, Sigma-Aldrich) was added for the last 4 h. After stimulation, cells were stained for flow cytometry as described below.

### Generation of bone marrow-derived dendritic cells

BMDCs were generated as described [95]. In brief, bone marrow was isolated from tibia and femur and filtered through a 70 µm strainer. Cells were differentiated in cRPMI medium supplemented with GM-CSF (from supernatant of GM-CSF-producing X63 cell line, concentration was pre-determined by titration) for 5 days at 37°C and 5% CO_2_. On days 2 and 3, the medium was changed and supplemented with fresh GM-CSF. On day 5, cells were collected and used for experiments.

### Stimulation of BMDCs

Samples of 10^5^ BMDCs per well were resuspended in cRPMI medium supplemented with GM-CSF and seeded in a 96-well plate. After around 2 h they were attached, and the stimuli were added. Fungal cultures were grown as explained above, washed, and diluted in cRPMI to a concentration of 10^6^ CFU/ml. Samples of100 μl per well were added to the BMDCs to receive a MOI of 1. Control stimuli Curdlan (100 μg/ml) and CPG (1000 ng/ml, Invivogen) were also resuspended in cRPMI and added to the BMDCs. Control samples of 100 μl cRPMI medium was used for unstimulated conditions.

### CLR-Fc staining for flow cytometry and microscopy

*Malassezia* spp. cultures were grown for 2 days as explained above, washed, and resuspended in PBS. The yeast cells were sonicated (10min; cycles of 15 sec bursts, 15 sec break) and filtered through a 40 µm strainer. Samples of 1×10^6^ cells were added to V-shaped 96 well plates and murine CLR-Fc’s: Dectin-1-Fc [26], Dectin-2-Fc [27], Mincle-Fc (Novus Biologicals) or CR-Fc [96] (negative control) [28] were added at 10 µg/ml. Following incubation at 4°C for 45 min, cells were washed, and bound CLR-Fc’s were detected with goat anti-human IgG Fc Alexa Fluor 488 (1:200, Thermo Fisher Scientific) for Dectin-1-Fc, Dectin-2-Fc and CR-Fc, or goat anti-mouse IgG Fc (1:200, Jackson ImmunoResearch) for Mincle-Fc. Following 30min at 4°C, unbound secondary antibody was washed away. For flow cytometry, cells were acquired on a BD Accuri C6 plus flow cytometer and analysed with FlowJo software. For microscopy, images were acquired using the Deltavision widefield microscope. Images were deconvoluted using the acquisition software and transferred to ImageJ for analysis.

### Flow cytometry

For analysis of Th17 cells, single cell suspensions of skin and dLN were stained with antibodies directed against surface antigens (Supplementary Table S1). LIVE/DEAD Fixable Near IR stain (Life Technologies) was used for exclusion of dead cells. After surface staining, murine cells were fixed and permeabilized using Cytofix/Cytoperm reagents (BD Biosciences) for subsequent intracellular staining with cytokine-specific antibodies diluted in Perm/Wash buffer (BD Bioscience, as appropriate. All staining steps were carried out on ice. Cells were acquired on a Spectral Analyzer SP6800 (Sony), a CytoFLEX S (Beckman Coulter) or a Cytek Aurora (Cytek) instrument. For analysis of myeloid cell dynamics in skin and dLN, we modified a published multi-color staining panel [35]. After incubation with Fc block (anti-CD16/32, 1:100, Clone S17011E, BioLegend), cells were stained with surface antibody mix (Supplementary Table S1), fixed and permeabilized using Cytofix/Cytoperm reagents, and then stained intracellularly. Cells were acquired on a Cytek Aurora instrument (Cytek). All data were analyzed with FlowJo software (FlowJo LLC). The gating of the flow cytometric data was performed according to the guidelines for the use of flow cytometry and cell sorting in immunological studies [97] [35], including pre-gating on viable and single cells for analysis. Absolute cell numbers were calculated based on a defined number of counting beads (BD Bioscience, Calibrite Beads) that were added to the samples before flowcytometric acquisition.

### Histology

Mouse tissue was fixed in 4 % (v/v) PBS-buffered paraformaldehyde overnight and embedded in paraffin. Sagittal sections (9µm) were stained with hematoxylin and eosin and mounted with Pertex (Biosystem, Switzerland) according to standard protocols. All images were acquired with a digital slide scanner (NanoZoomer 2.0-HT, Hamamatsu) and analyzed with NDP view2 (Hamamatsu).

### RNA extraction and RT-qPCR

Isolation of total RNA from snap-frozen BMDCs was performed using TRI reagent (Sigma-Aldrich) according to the manufacturer’s instructions. cDNA was generated by RevertAid reverse transcriptase (ThermoFisher) and random nonamer oligonucleotides. Quantitative PCR was performed using SYBR green (Roche) and a QuantStudio 7 Flex instrument (Life Technologies). The primers used for qPCR were *Actb* forward 5’-CCCTGAAGTACCCCATTGAAC-3’, *Actb* reverse 5’-CTTTTCACGGTTGGCCTTAG-3’; *Il23a* forward 5’- CCAGCAGCTCTCTCGGAATC-3’, *Il23a* reverse 5’- TCATATGTCCCGCTGGTGC- 3‘; *Il6* forward 5’- GAGGATACCACTCCCAACAGACC-3’, *Il6 reverse* 5’-AAGTGCATCATCGTTGTTCATACA-3‘. All qPCR reactions were performed in duplicates, and the relative expression (rel. expr.) of each gene was determined after normalization to *Actb* transcript levels.

### Cytokine quantification by ELISA

IL12/IL23p40 levels in the supernatant of stimulated BMDCs were quantified using anti-mouse IL-12/IL-23p40 (clone C15.6, Thermo Fisher Scientific) for coating and biotinylated anti-mouse IL-12/IL-23p40 (clone C17.8, Thermo Fisher Scientific) in combination with ExtrAvidin®-Alkaline Phosphatase (Sigma) for detection. For the standard, recombinant mouse IL-12 (Biosource) was used.

### RNA-Sequencing data analysis

For RNA sequencing analysis we explored a published data set (NCBI GEO repository, accession number GSE253214 [22]. Data analysis was performed using the SUSHI framework [98], including differential expression using the generalized linear model as implemented by the DESeq2 Bioconductor R package [99], and Gene Ontology (GO) term pathway analysis using the hypergeometric over-representation and GSEA tests via the ‘enrichGO’ and ‘gseG’ functions respectively of the clusterProfiler Bioconductor R package [100]. Figures were generated using the exploreDEG Interactive Shiny App (https://doi.org/10.5281/zenodo.8167438). All R functions were executed on R version 4.1 (R Core Team, 2020) and Bioconductor version 3.14.

## DATA AVAILABILITY STATEMENT

All raw data linked to this study will be made publicly available at zenodo.org upon acceptance of the manuscript (doi will be provided).

## ACKNOWLEDGEMENTS (INCLUDING FUNDING)

The authors would like to thank Gordon Brown, Thorsten Buch, Manfred Kopf, Mark Suter, Marc Vocanson and Wolf-Dietrich Hardt for mice; David Sancho for bone marrow; the staff of the Laboratory Animal Service Center of University of Zurich for animal husbandry; the staff of the Laboratory for Animal Model Pathology of University of Zurich for histology and members of the LeibundGut-lab for helpful advice and discussions. This work was supported by the Swiss National Science Foundation (grant #310030_189255 to SLL) and the NIH (grant #1R21-AI168672-01A1 to SLL). NARG acknowledges support of Wellcome Trust Investigator, Collaborative, Equipment, Strategic and Biomedical Resource awards (101873, 200208, 215599, 224323). NARG and MHTS thank the MRC (MR/M026663/2), the MRC Centre for Medical Mycology (MR/N006364/2) and the National Institute for Health and Care Research (NIHR) Exeter Biomedical Research Centre (BRC). The funders had no role in study design, data collection and analysis, decision to publish, or preparation of the manuscript.

## AUTHOR CONTRIBUTIONS

Conceptualization: Meret Tuor, Salomé LeibundGut-Landmann.

Formal analysis: Meret Tuor, Mark Stappers, Alice Desgardin.

Funding acquisition: Salomé LeibundGut-Landmann.

Investigation: Meret Tuor (all figures except for 1B-D, 4A-B, S1A-B, S4A-B), Mark Stappers (Fig. 1B-D, Fig. S1A-B), Fiorella Ruchti (Fig. 4A-B), Alice Desgardin (Fig. 4A-B), Florian Sparber (Fig. S4A-B).

Resources: Selinda Orr

Project administration: Salomé LeibundGut-Landmann, Neil Gow.

Supervision: Salomé LeibundGut-Landmann, Neil Gow.

Validation: Salomé LeibundGut-Landmann, Neil Gow.

Visualization: Meret Tuor.

Writing – original draft: Meret Tuor, Salomé LeibundGut-Landmann.

Writing – review & editing: all authors

## COMPETING INTERESTS

The authors declare that they have no conflicts of interest to disclose.

## Abbreviations (abbreviations that are used three or more times in the text)

APCs: antigen- presenting cells
BMDCs: bone marrow-derived dendritic cells
CLR: C-type lectin receptor
DC: dendritic cell
dLN: draining lymph nodes
IL-17: interleukin-17
LCs: Langerhans cells
MFI: median fluorescence intensity
PRR: pattern recognition receptor
T helper 17 cells: Th17
TLR: toll-like receptor
DKO: double knockout
TKO: triple knockout

## SUPPLEMENTARY FIGURE LEGENDS

**Figure S1 (Related to Figure 1). The C-type lectin receptors Mincle, Dectin-1, and Dectin-2 bind to *Malassezia* spp.**

Live *M. pachydermatis*, *M. sympodialis* and *M. furfur* cells were incubated with Mincle-Fc, Dectin-1-Fc, Dectin-2-Fc or controls (2nd (amIgG) or CR-Fc) and analyzed by flow cytometry. **A.** Gating strategy for single yeast cells among cell events.

**Figure S2 (Related to Figure 2). CLR-Card9-dependent signaling in response to *Malassezia* activates dendritic cells.**

**A-C.** BMDCs were generated from *Card9*^-/-^ and *Card9*^+/-^ mice and stimulated with curdlan or CpG (A), or with live *M. sympodialis* or *M. furfur* for 24h (B, C). IL-12/IL-23p40 secretion was quantified by sandwich ELISA n = 9,9 (A), *Il23a* and *Il6* expression was quantified by RT-qPCR, n = 3, 3, one independent experiment, mean ± SD (B, C). **D-G.** BMDCs generated from *Clec4e*^-/-^ and WT control mice (D), *Clec7a*^-/-^ and *Clec7a*^+/-^ mice (E), *Clec4n*^-/-^ and *Clec4n*^+/-^ mice (F), and Dectin-1-Dectin-2 DKO, Mincle-Dectin-2-Dectin-1 TKO and WT control mice (G) were stimulated with curdlan or CpG for 24 h. IL-12/IL-23p40 secretion was quantified by sandwich ELISA. C: n = 6, 6, pooled from two independent experiments, mean ± SEM; D: n = 6, 9, pooled from three independent experiments, mean ± SEM; E: n = 6, 6, pooled from two independent experiments, mean ± SEM; F: n = 6, 6, 6, pooled from two independent experiments, mean ± SEM. Statistical significance was determined using, unpaired t test. In A, D-G, the t test was conducted for each stimulus separately, * p < 0.05, *** p<0.001, **** <0.0001.

**Figure S3 (Related to Figure 3). Gating strategy for myeloid cells in the ear skin and dLN.**

**A-B.** Gating strategy for myeloid cell types in ear skin (A) and dLN (B) shown in Figure 3. **C.** Representative histogram of CD86 expression by dLN cDC2 cells 2 and 3 days after *M. pachydermatis* colonization. **D.** Representative plots showing IL-12/IL-23p40^+^ cDC2 cells 2 and 3 days after *M. pachydermatis* colonization. Data in C and D are from one representative of three independent experiments with n= 5, 4, 5.

**Figure S4 (Related to Figure 3). *Malassezia* skin colonization recruits myeloid cells and activates cDC2s.**

**A-B.** The ear skin of *Batf3*^+/+^ and *Batf3*^-/-^ mice was colonized with *M. pachydermatis* for 6 days. XCR1^+^ cDC1 cells within CD11c^+^ MHC-II^+^ DCs were quantified in the ear skin (A). CD4^+^ T cell counts, IL-17A^+^ CD4^+^ CD44^+^ T cell counts, and the % of IL-17A producing CD4^+^ CD44^+^ T cells were quantified in the dLN (B). n = 13, 4, data from one representative of two independent experiments, mean ± SD. **C-F.** Langerin-DTR mice were treated with 1 μg DT or PBS control (-DT) one day before and on the day of fungal colonization and cell recruitment was assessed after 7 days. EpCAM^+^ LCs within CD11c^+^ MHC-II^+^ Sirpα^+^ DCs were quantified in ear skin (C) or dLN (D). CD4^+^ T cell counts, IL-17A^+^ CD4^+^ CD44^+^ T cell counts, and the % of IL- 17A producing CD4^+^ CD44^+^ T cells were quantified in ear skin (E) and dLN (F). n = 7, 6, data pooled from two independent experiments, mean ± SEM. Statistical significance was determined using unpaired t test, * p < 0.05, ** <0.01, **** p < 0.0001.

**Figure S5 (Related to Figure 4). Dectin-2 but not Mincle nor Dectin-1 mediates Th17 immunity to *Malassezia*.**

**A.** Gating strategy for identifying CD4^+^ cells among CD90^+^ cells, and IL-17A-producing, IL-22- producing and IL-17A, IL-22 double producing cells among CD4^+^CD44^+^ T cells. **B-C.** Quantification of IL-22 and IL-17A, IL-22 double producing CD4^+^ T cells in ear skin (B) or dLN (C) of *Card9*^-/-^ and *Card9*^+/-^ mice, colonized with *M. pachydermatis* for 7 days. n = 8, 8, data pooled from two independent experiments, mean ± SEM. **D-E.** Quantification of CD4^+^ T cell counts, IL-17A^+^ CD4^+^ CD44^+^ T cell counts, and the % of IL-17A producing CD4^+^ CD44^+^ T cells in ear skin (D) and dLN (E) of non-colonized *Card9*^-/-^ and *Card9*^+/-^ mice. n = 10, 10, data pooled from two independent experiments, mean ± SEM. **F-H.** Mincle chimeric mice. CD45.1+ host and CD45.2+ donor cells among all live cells in the dLN of chimeric mice 6-8 weeks after reconstitution (F). Quantification of IL-22 and IL-17A, IL-22 double producing CD4^+^ CD44^+^ T cells in ear skin (G) or dLN (H). n = 8, 8, data pooled from three independent experiments, mean ± SEM. **I-J.** Quantification of IL-22 and IL-17A, IL-22 double producing CD4^+^ T cells in ear skin (I) or dLN (J) of *Clec7a*^-/-^ (Dectin-1) and *Clec7a*^+/-^ mice. n = 10, 9, data pooled from two independent experiments, mean ± SEM. Statistical significance was determined using unpaired t test, * p < 0.05, ** p < 0.01.

**Figure S6 (Related to Figure 4). Dectin-2 but not Mincle nor Dectin-1 mediates Th17 immunity to *Malassezia*.**

**A-B.** Quantification of IL-22 and IL-17A, IL-22 double producing CD4^+^ T cells in ear skin (A) or dLN (B) of *Clec4n*^-/-^ and *Clec4n*^+/-^ female mice colonized with *M. pachydermatis* for 7 days. n = 6, 6, data pooled from two independent experiments, mean ± SEM. **C-D.** Quantification of CD4^+^ T cell counts, IL-17A^+^ CD4^+^ CD44^+^ T cells in ear skin (C) and dLN (D) in *Clec4n*^-/-^ and *Clec4n*^+/-^ male mice. n = 6, 6, data pooled from two independent experiments, mean ± SEM. **E-H.** The ear skin of *Clec4n*^-/-^ and *Clec4n*^+/-^ mice (E, F) and *Clec7a*^-/-^ and *Clec7a*^+/-^ mice (G, H) was colonized with *M. sympodialis* for 7 days. Quantification of CD4^+^ T cell counts, IL-17A^+^ CD4^+^ CD44^+^ T cell counts, and the % of IL-17A producing CD4^+^ CD44^+^ T cells in ear skin (E, G) and dLN (F, H). E, F: n = 3, 4, female mice only, data from one representative experiment, mean ± SD G, H: n = 10, 9, data pooled from two independent experiments, mean ± SEM. **I-J.** Quantification of CD4^+^ T cell counts, IL-17A^+^ CD4^+^ CD44^+^ T cell counts, and the % of IL-17A producing CD4^+^ CD44^+^ T cells in ear skin (E) and dLN (F) of non-colonized *Clec4n*^-/-^ and *Clec4n*^+/-^ mice. n = 4, 5, data from one representative experiment, mean ± SD. **K.** DKO and TKO chimeric mice. CD45.1+ host and CD45.2+ donor cells among all live cells in the dLN 6-8 weeks after reconstitution. Statistical significance was determined using unpaired t test, * p < 0.05, ** p < 0.01.

**Figure S7 (Related to Figure 5). The Th17 response against *Malassezia* depends on T-cell-intrinsic MyD88 signaling.**

**A.-B.** BMDCs were generated from *Tlr23479* ^-/-^ and WT mice (A) or *MyD88*^-/-^ and *MyD88*^+/-^ mice (B) and stimulated with curdlan and CpG controls or with live *M. pachydermatis*, *M. sympodialis* or *M. furfur* for 24 h. IL-12/IL-23p40 secretion was quantified by sandwich ELISA. Each symbol represents an independently stimulated well. The mean ± SEM or SD is indicated for each group. Data are pooled from two independent experiments (A) or are from a single experiment (B). **C-D.** Quantification of IL-22 and IL-17A, IL-22 double producing CD4^+^ T cells in ear skin (C) or dLN (D) of *MyD88*^-/-^ and *MyD88*^+/-^ mice 7 days after *M. pachydermatis* colonization. n = 10, 11, data pooled from three independent experiments, mean ± SEM. **E-F.** Quantification of CD4^+^ T cell counts, IL-17A^+^ CD4^+^ CD44^+^ T cell counts, and the % of IL-17A producing CD4^+^ CD44^+^ T cells in ear skin (E) and dLN (F) of non-colonized *MyD88*^-/-^ and *MyD88*^+/-^ mice. n = 4, 4, data from one experiment, mean ± SD. **G.** CD45.1+ and CD45.2+ host and donor cells among all live cells in the dLN of chimeric mice from Fig. 5E-F 6-8 weeks after reconstitution and 7 days after *M. pachydermatis* colonization. **H.** The ratio of CD45.1^+^ and CD45.2^+^ cells in mixed chimeras from Figure 5G-H at the time point of reconstitution (left) and 7 days after *M. pachydermatis* colonization in 6-8 week- reconstituted chimeras (right). Statistical significance was determined using, unpaired t test (A-F). For curdlan and CpG stimulations, the t test was conducted for each stimulus separately, * p < 0.05, ** p < 0.01, *** p<0.001, **** <0.0001.

**Figure S8 (Related to Figure 6). *Malassezia*-induced inflammatory IL-1 signaling induces Th17 responses.**

**A-B.** Quantification of IL-22 producing CD4^+^ T cells in ear skin (A) or dLN (B) of *Caspase1* ^-/-^ and WT mice. n = 10, 10, data pooled from two independent experiments, mean ± SEM. **C-D.** Quantification of IL-22 and IL-17A, IL-22 double producing CD4^+^ T cells in ear skin (C) or dLN (D) of *Il1rl2*^-/-^ and *Il1rl2*^+/-^ mice. n = 10, 10, data pooled from two independent experiments, mean ± SEM. Statistical significance was determined using unpaired t test, * p < 0.05, ** p < 0.01.

## Notes

### Competing Interest Statement

The authors have declared no competing interest.

