## Supplementary figures and images for "Card9 and MyD88 differentially regulate Th17 immunity to the commensal yeast *Malassezia* in the murine skin"

### Supplementary Figures S1 - S8

**Fig S1**

**A**

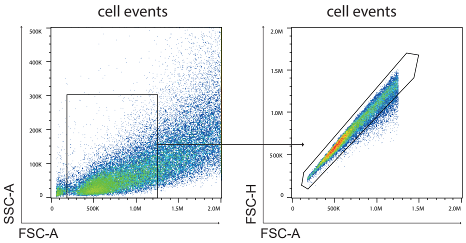

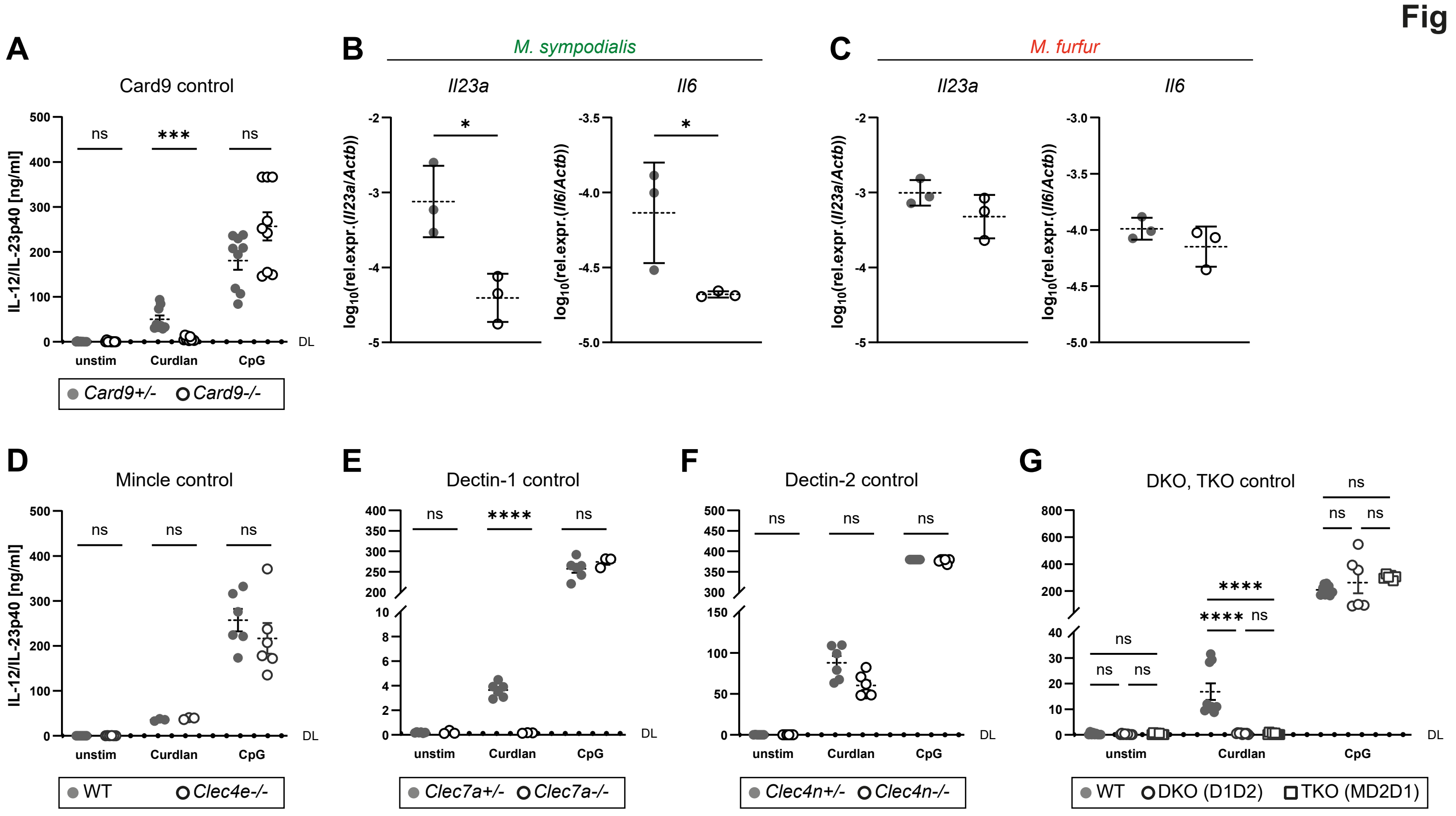

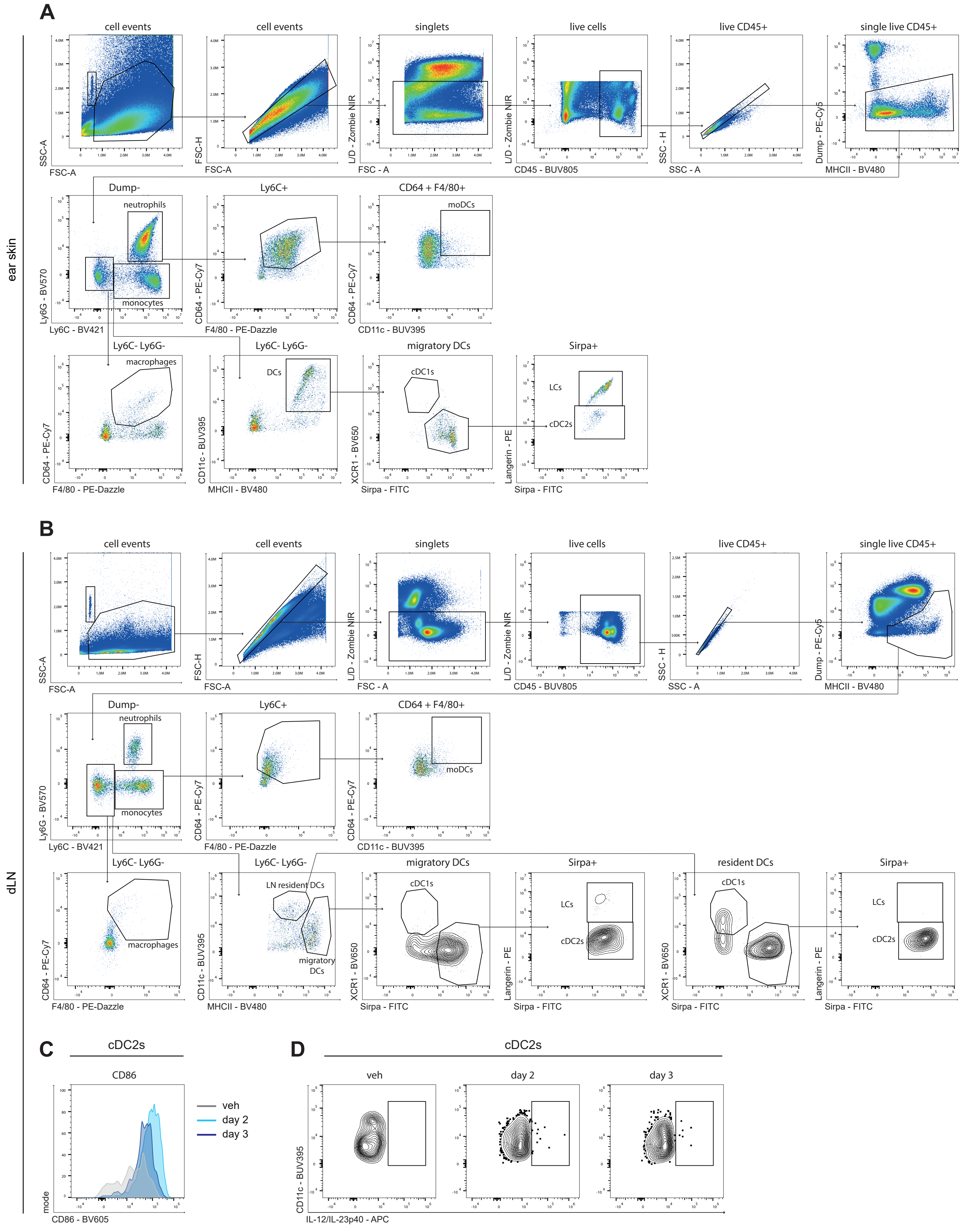

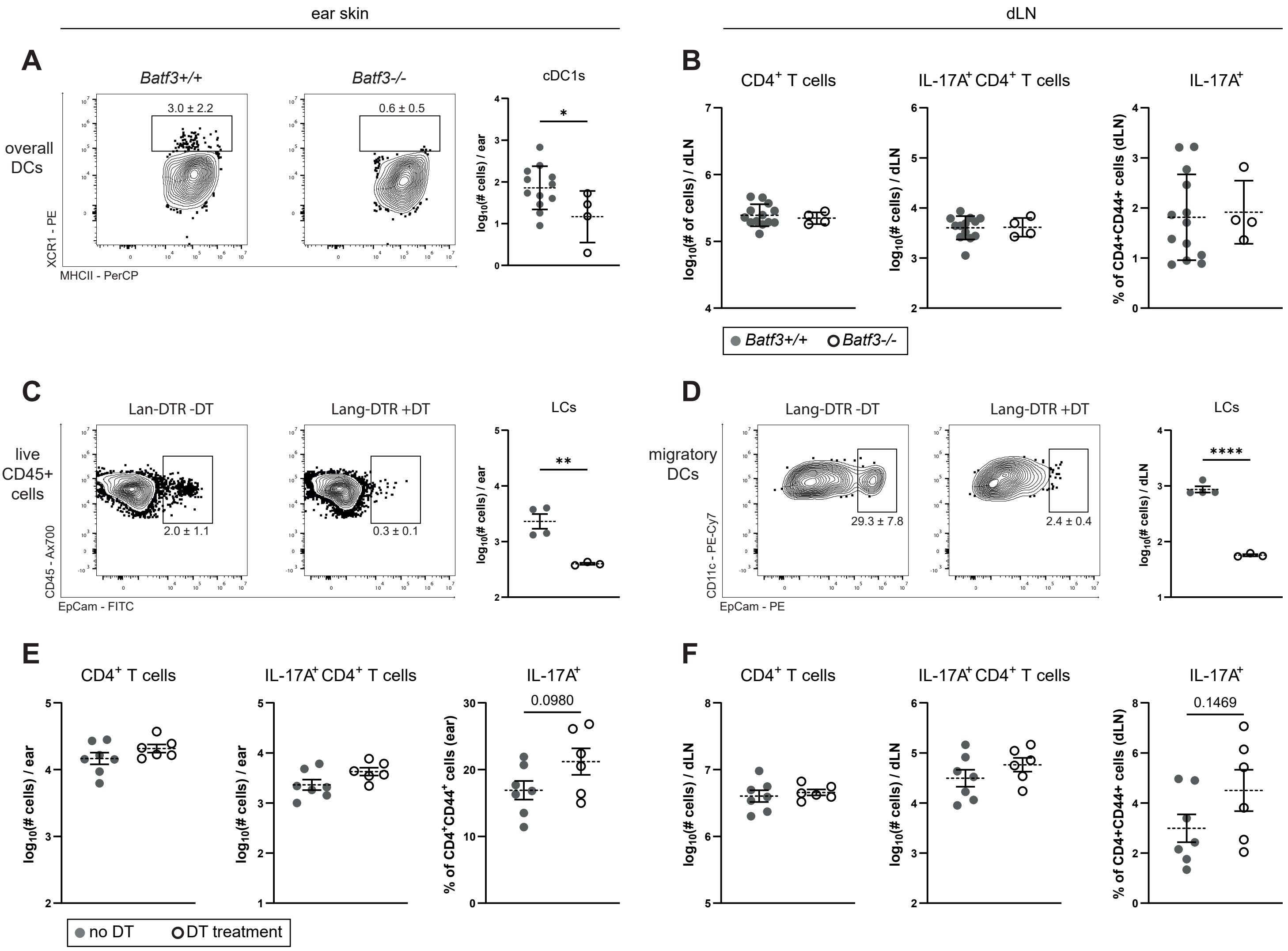

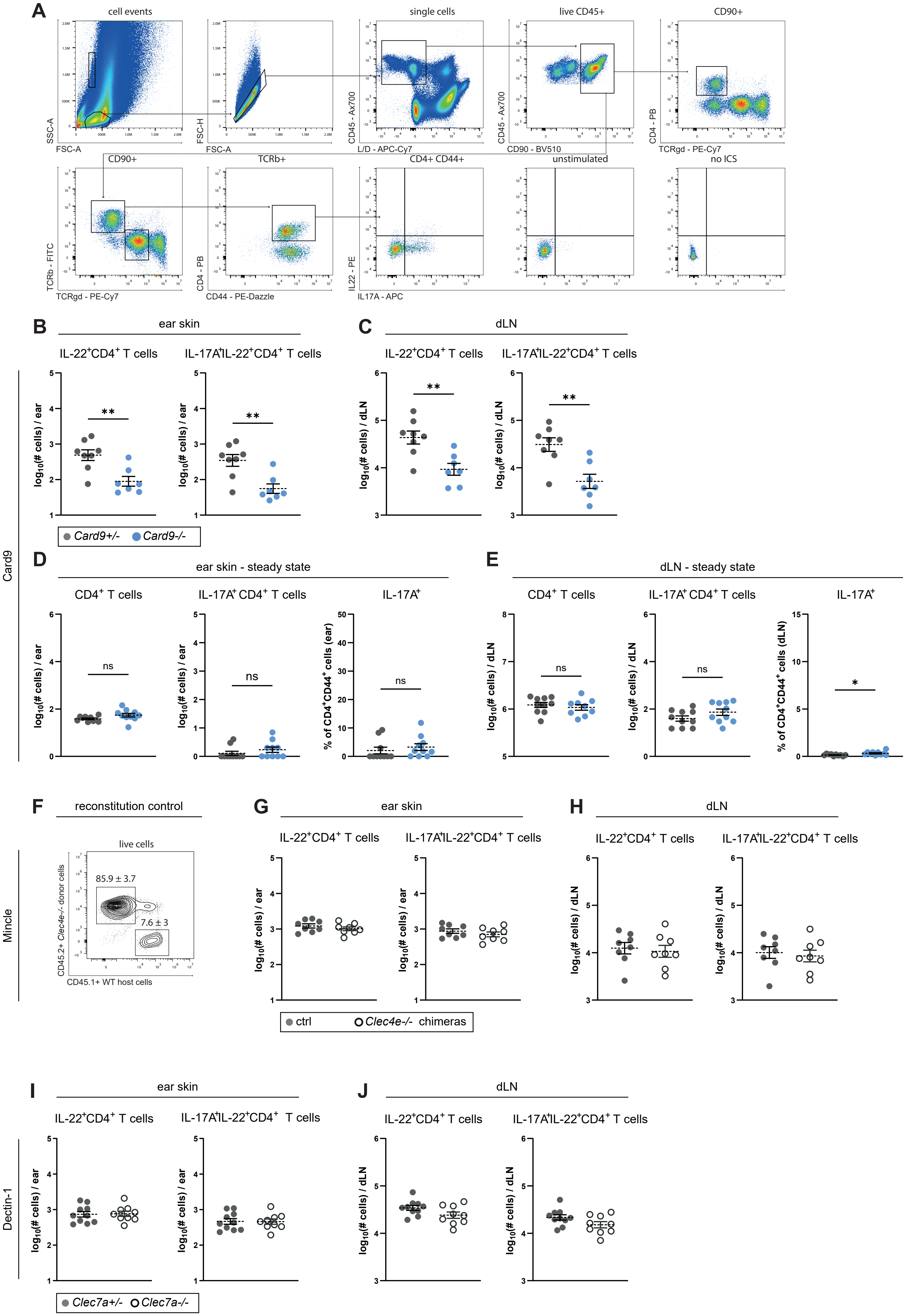

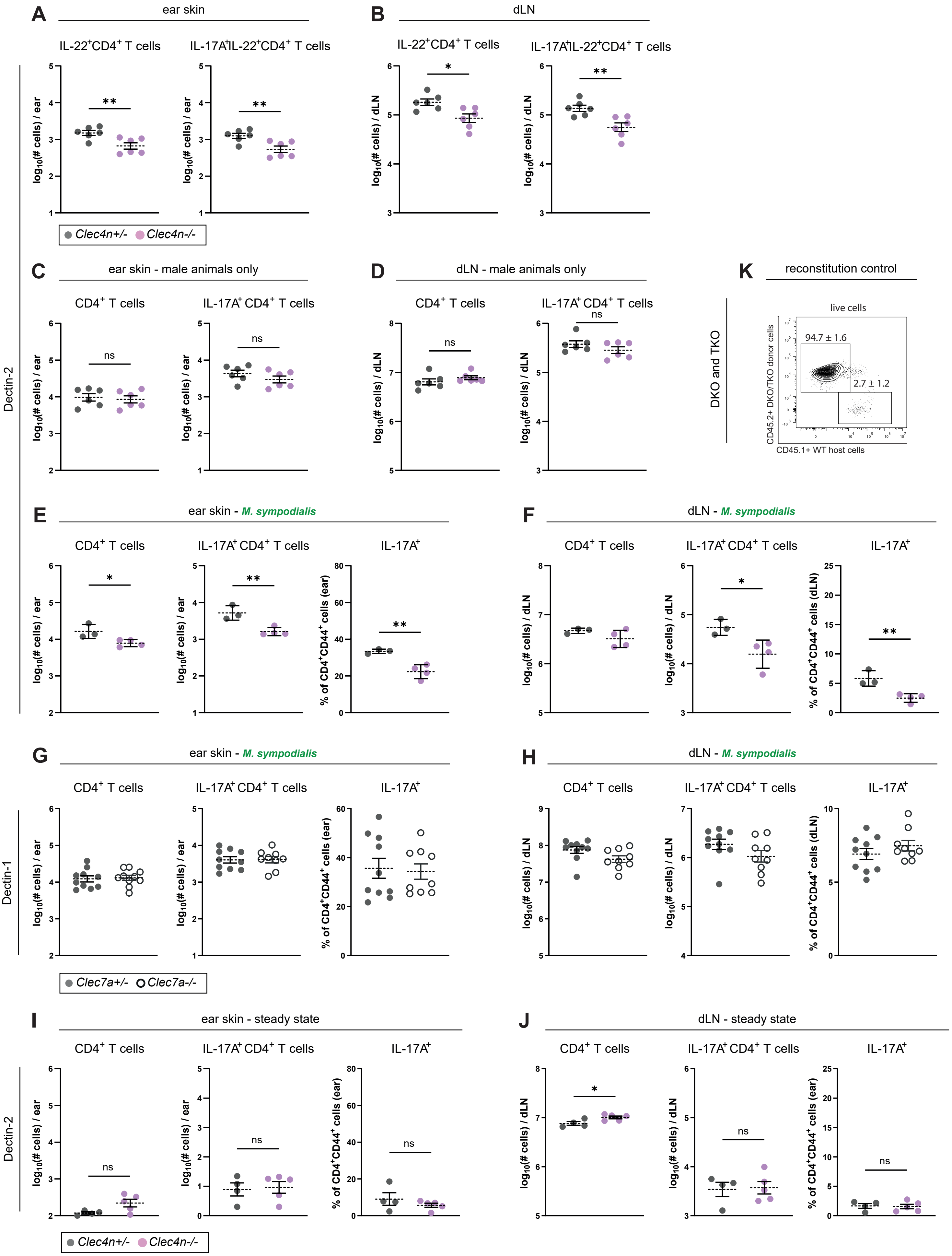

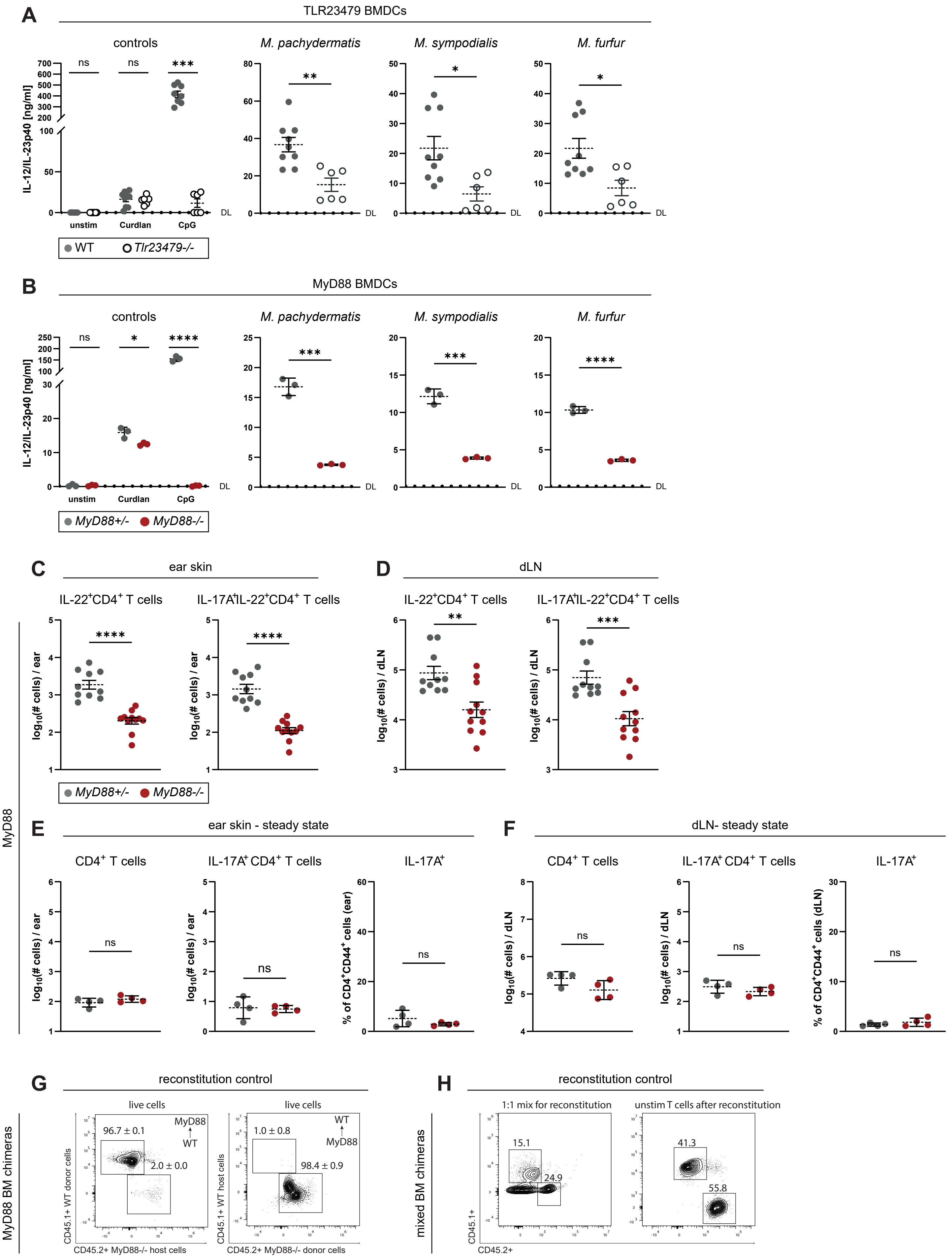

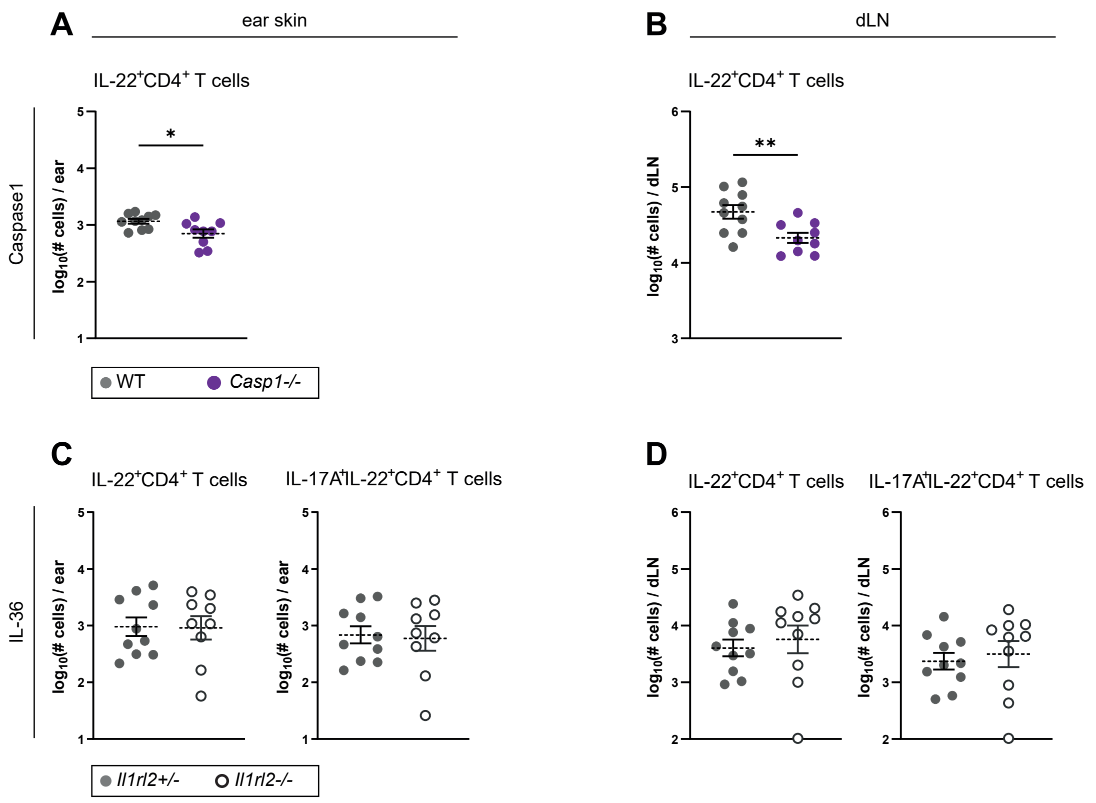
