## Supplementary Table S1 for "Card9 and MyD88 differentially regulate Th17 immunity to the commensal yeast *Malassezia* in the murine skin"

| **Antibodies (clones)** | **Source** | **Identifier** |
| --- | --- | --- |
| **Anti-mouse** | | |
| FITC anti-CD11b (M1/70) | Biolegend | Cat#101205, RRID:AB_312788 |
| FITC anti-CD19 (6D5) | Biolegend | Cat#115506, RRID:AB_313641 |
| FITC anti-CD4 (RM4-4) | Biolegend | Cat#116003, RRID:AB_313688 |
| FITC anti-CD44 (IM7) | Biolegend | Cat#103022, RRID:AB_493685 |
| FITC anti-CD8α (53-6.7) | Biolegend | Cat#100706, RRID:AB_312745 |
| FITC anti-CD90.2 (30-H12) | Biolegend | Cat#105305, RRID:AB_313176 |
| FITC anti-Ly6G (1A8) | Biolegend | Cat#127605, RRID:AB_1236488 |
| FITC anti-MHCII (M5/114.15.2) | Biolegend | Cat#107605, RRID:AB_313320 |
| FITC anti-TCRβ (H57-597) | Biolegend | Cat#109205, RRID:AB_313428 |
| FITC anti-Vγ4 (Vγ2) (UC3-10A6) | Biolegend | Cat#137704, RRID:AB_10569353 |
| PE anti-CCR2 (SA203G11) | Biolegend | Cat#150609, RRID:AB_2616981 |
| PE anti-IL-17A (TC11-18H10.1) | Biolegend | Cat#506904, RRID:AB_315464 |
| PE anti-Vγ4 (Vγ2) (UC3-10A6) | Biolegend | Cat#137705, RRID:AB_10643997 |
| PE-Dazzle anti-CD27 (LG.3A10) | Biolegend | Cat#124227, RRID:AB_2565793 |
| PE-Dazzle anti-CD44 (IM7) | Biolegend | Cat#103055, RRID:AB_2564043 |
| PE-Dazzle anti-CXCR6 (SA051D1) | Biolegend | Cat#151116, RRID:AB_2721699 |
| PE-Dazzle anti-CD103 (2E7) | Biolegend | Cat#121429, RRID:AB_2566492 |
| PerCP anti-MHCII (M5/114.15.2) | Biolegend | Cat#107624, RRID:AB_2191073 |
| PE-Cy5 anti-CD3ε (145-2C11) | Biolegend | Cat#100310, RRID:AB_312675 |
| PE-Cy7 anti-IL-17A (TC11-18H10.1) | Biolegend | Cat#506922, RRID:AB_2125010 |
| PE-Cy7 anti-TCRγδ (GL3) | Biolegend | Cat#118124, RRID:AB_11204423 |
| Pacific Blue anti-CD103 (2E7) | Biolegend | Cat#121418, RRID:AB_2128619 |
| Pacific Blue anti-CD4 (RM4-5) | Biolegend | Cat#100531, RRID:AB_493374 |
| Pacific Blue anti-Ly6G (1A8) | Biolegend | Cat#127612, RRID:AB_2251161 |
| BV421 anti-CCR6 (29-2L17) | Biolegend | Cat#129817, RRID:AB_10898320 |
| BV510 anti-CD90 (30-H12) | Biolegend | Cat#105335, RRID:AB_2566587 |
| BV570 anti-CD90 (30-H12) | Biolegend | Cat#105329, RRID:AB_10917055 |
| BV605 anti-CD11b (M1/70) | Biolegend | Cat#101237, RRID:AB_11126744 |
| BV605 anti-CD4 (RM4-4) | Biolegend | Cat#116027, RRID:AB_2800581 |
| APC anti-CD11c (N418) | Biolegend | Cat#117310, RRID:AB_313779 |
| APC anti-CD44 (IM7) | Biolegend | Cat#103012, RRID:AB_312963 |
| APC anti-IL-17A (TC11-18H10.1) | Biolegend | Cat#506916, RRID:AB_536018 |
| AF700 anti-Ki-67 (16A8) | Biolegend | Cat#652419, RRID:AB_2564284 |
| AF700 Anti-CD45.2 (104) | Biolegend | Cat#109821, RRID:AB_493730 |
